# LRRC8A-containing anion channels promote glioblastoma proliferation via a WNK1/mTORC2-dependent mechanism

**DOI:** 10.1101/2025.02.02.636139

**Authors:** Antonio M. Fidaleo, Martin D. Bach, Shaina Orbeta, Iskandar F. Abdullaev, Nina Martino, Alejandro P. Adam, Mateo A. Boulos, Nickolai O. Dulin, Alexandra R. Paul, Yu-Hung Kuo, Alexander A. Mongin

**Author notes:** These authors contributed equally to this work and therefore should be considered co-first authors (Antonio M. Fidaleo, Martin D. Bach).

## Abstract

Leucine-rich repeat-containing protein 8A (LRRC8A) is the essential subunit of ubiquitous volume-regulated anion channels (VRACs). LRCC8A is overexpressed in several cancers and promotes negative survival outcomes via a poorly defined mechanism. Here, we explored the role of LRRC8A and VRACs in the progression of glioblastoma (GBM), the most common and deadly primary brain tumor. We found that, as compared to healthy controls, *LRRC8A* mRNA was strongly upregulated in surgical GBM specimens, patient-derived GBM cell lines, and GBM datasets from The Cancer Genome Atlas (TCGA). Our in-silico analysis indicated that patients belonging to the lowest *LRRC8A* expression quartile demonstrated a trend for extended life expectancy. In patient-derived GBM cultures, siRNA-driven LRRC8A knockdown reduced cell proliferation and additionally decreased intracellular chloride levels and inhibited activity of mTOR complex 2. The antiproliferative effect of LRRC8A downregulation was recapitulated with a pharmacological inhibitor of VRAC. Our ensuing biochemical and molecular biology analyses established that the LRRC8A-containing VRACs facilitate GBM proliferation via a new mechanism involving non-enzymatic actions of the chloride-sensitive protein kinase WNK1. Accordingly, the chloride-bound WNK1 stimulates mTORC2 and the mTORC2-dependent protein kinases AKT and SGK, which promote proliferation. These findings establish the new mTORC2-centric axis for VRAC dependent regulation of cellular functions and uncover potential targets for GBM intervention.

**SUBJECT AREAS:** Cell biology

Molecular biology

Cancer

## INTRODUCTION

Glioblastoma (GBM) is the most common and aggressive primary brain cancer in adults that arises due to the accumulation of somatic mutations in neural or glial precursor cells.^1,2,3^ With an annual incidence rate of ∼3.2 cases per 100,000 people, GBM accounts for approximately half of all primary brain malignancies.^4^ Compared to other brain tumors, GBM is associated with disproportionately high morbidity and mortality, with the majority of patients dying within two years of diagnosis and only 5-6% surviving past five years.^4,5^ Based on extensive clinical evidence, the standard of care for GBM patients includes maximal safe tumor resection followed by radio- and chemotherapy with the brain-penetrable alkylating agent temozolomide.^6,7,8^ Unfortunately, the highly infiltrative nature of GBM makes complete removal nearly impossible; furthermore, GBM tumors rapidly develop resistance to the first-line therapies and invariantly recur. ^5,7,8^ Even the most aggressive treatment regiments increase median survival time only marginally and do not extend median life expectancy past 15-16 months.^5,7^ The inherent resistance of GBMs to all current treatment modalities, alongside the high degree of intertumoral and intratumoral heterogeneity, necessitates the search for new prospective therapeutic targets. Some of the recently developed and tested approaches include tumor-treating fields^9^, viral and immune therapies (e.g.,^10,11,12^), and inhibitors of signaling pathways associated with GBM progression (e.g.,^13,14^).

Cancer development is frequently associated with profound changes in the expression and/or activity of ion channels, leading to the idea that cancerogenesis may be viewed as an ‘onco-channelopathy’ (see^15,16^, and overview in^17^). Modified ion channel function has been linked to many cancer hallmarks, including unrestricted proliferation, tissue invasion, metastasis, pathological angiogenesis, and evasion of apoptosis (reviewed in^15,16,18,19,20,21,22^). Chloride (Cl^−^) channels are a diverse group of membrane proteins which facilitate movement of Cl^−^, bicarbonate, and various small organic molecules. They regulate numerous cellular processes, including many of those relevant to cancer biology. ^23,24,25^ Among many distinct Cl^−^ channels, volume regulated anion channels (VRACs) have long been speculated to contribute to cancer progression^26,27^, including in GBM^28,29,30^. The ubiquitous VRACs, also known as volume-sensitive organic osmolyte-anion channels (VSOAC) or volume-sensitive outward rectifying chloride channels (VSOR), are activated by cell swelling and are the key players in cell volume regulation.^26,31,32^. However, they are also thought to be important for cell proliferation, migration, angiogenesis, apoptotic cell death, and chemoresistance.^22,26,33,34,35^

For a long time, progress in VRAC research was impeded by a lack of knowledge regarding the channel’s molecular composition.^36^ This problem persisted until about ten years ago, when two seminal studies discovered that VRACs are hexameric complexes composed of proteins from the leucine-rich repeat-containing family 8 (LRRC8), with one of these, LRRC8A, playing an indispensable role in channel function.^37,38^ These new findings rapidly revolutionized the field and dramatically expanded our understanding of the physiological importance of VRACs. Thus, it was demonstrated that LRRC8A can serve as both a subunit of VRAC and as a signaling scaffold for several proteins linked to growth factor signaling.^39,40,41,42^ Using a conditional deletion of the essential LRRC8A, VRACs were shown to contribute to many diverse physiological processes such as glucose sensing and metabolic regulation, fertility, paracrine signaling, immune function, and others.^40,43,44,45^ Among important recent developments, several laboratories discovered that LRRC8A is upregulated in a variety of cancers and that its expression levels are inversely related to patients’ life expectancy (e.g.,^46,47,48,49^). Our group was the first to find that LRRC8A-containing VRACs regulate GBM proliferation and sensitivity to the chemotherapeutic agent temozolomide.^50^ Here, we performed an extensive analysis of LRRC8A expression and function in human GBM and explored the intracellular mechanisms linking VRAC activity to enhanced proliferation in patient-derived GBM cells. We discovered a previously unknown Cl^−^ dependent mTOR complex 2-centric mechanism linking VRAC activity to cancer cell proliferation.

## RESULTS

### LRRC8A expression is upregulated in GBM tumor tissue and in patient-derived GBM cells

Based on our previous discovery of the potentially important role of LRRC8A in GBM proliferation^50^, we analyzed expression levels for each of the five LRRC8 family members in surgically resected GBM specimens. We employed a qRT-PCR approach to compare 22 GBM samples to three separate brain tissue RNA medleys isolated postmortem from healthy brains and representing a total of 8 individuals (see *Star*⋆*Methods* for details). The main finding of this comparison was the dramatic upregulation of LRRC8A expression in the majority of GBM specimens which, on average, was six-times higher than in non-pathological brain tissue (Figure 1A1, p<0.001). In contrast, LRRC8B, LRRC8C, and LRRC8D gene products were present in comparable quantities in both malignant and non-malignant brain samples (Figure 1A1). Consistent with other reports and prior RNA-seq analyses of brain cells (see^51^), LRRC8E expression was either extremely low or undetectable (not shown). The abundance of LRRC8A mRNA in GBM was much higher than that of other LRRC8 isoforms, which is unusual when compared to the LRRC8 expression patterns in non-malignant tissues (see *Discussion*).

Keeping in mind the heterogeneity of tumor material, which contains vasculature and large numbers of invading immune cells, we extended qRT-PCR analysis of LRRC8/VRAC expression to primary GBM cell cultures derived from the surgical specimens. In these experiments, expression was compared to primary human astrocytes cultivated under identical conditions. Similar to GBM tissue data, we found that LRRC8A mRNA expression was significantly elevated, approximately five-fold, as compared to non-malignant astrocytes (Figure 1A2, p<0.05). Interestingly, we found that GBM cultures also had higher expression of LRRC8B, LRRC8C and LRRC8D subunits (Figure 1A2, adjusted p<0.05 for all transcripts). Still, the pattern of overabundance of LRRC8A mRNA relative to the other members of LRRC8 family strongly resembled that seen in surgical GBM specimens.

For independent validation of our findings, we evaluated clinical RNA-seq data available through The Cancer Genome Atlas (TCGA) program. We selected the TCGA GBM datasets that included full RNA-seq and survival information and were confirmed as expressing wild type IDH1/2 transcript. The patients with mutated IDH1/2 were excluded, as this mutation is indicative of the biologically distinct secondary GBM.^52^ The transcript abundance of LRRC8 isoforms was compared between 126 primary GBMs and 3 RNA medleys representing 29 healthy brains. The healthy brain data were sourced from the Gene Expression Omnibus repository (GEO series GSE196695, see^53^). In our in-silico RNA-seq analysis, the pattern of LRRC8 subunit expression largely mirrored what was seen in our internally obtained GBM specimens. LRRC8A was by far the most prominently expressed subunit, with the median expression in GBM samples approximately 3-fold higher than in healthy brains (Figure 1A3, p<0.001). LRRC8B through D were expressed at much lower levels than LRRC8A, in quantities comparable across GBM and healthy brain tissue (Figure 1A3). LRRC8E was consistently low-to-undetectable throughout all datasets (not shown).

Overall, the pattern of elevated expression of LRRC8A in GBM was remarkably similar in all three analyses. In contrast, LRRC8B-D subunits were expressed in comparable quantities in malignant tissue and healthy brains.

### LRRC8A-containing VRACs are expressed in patient-derived GBM cells and are important for GBM proliferation

Next, we assessed the functional expression and physiological relevance of LRRC8A-containing VRACs in patient-derived GBM cells. To account for intertumoral heterogeneity, we performed our in-vitro analyses in two different GBM cell lines, here referred to as GBM1 and GBM8. Both were isolated and propagated from clinically diagnosed GBM specimens but had distinct morphological and molecular features. GBM1 cells had a dedifferentiated morphology (Figure 1B and Figure S1A), very high proliferative capacity, and low-to-undetectable GFAP expression (Figure S1C). In contrast, GBM8 had an epithelial morphology resembling that of cultured astrocytes (Figure 1B and Figure S1B), lower rate of proliferation, and high expression of the astrocytic marker GFAP (Figure S1C). Consistent with their astroglial lineage, GBM1 and GBM8 had high glutamate uptake rates comparable to human astrocytes (Figure S1E-G). qRT-PCR analysis revealed moderate differences in the levels of LRRC8A mRNA expression between two primary cell lines. In GBM1, LRRC8A mRNA levels were similar to those in human astrocytes, while GBM8 had 2-4-fold higher LRRC8A mRNA abundance (Figure S1D).

**Figure 1.**
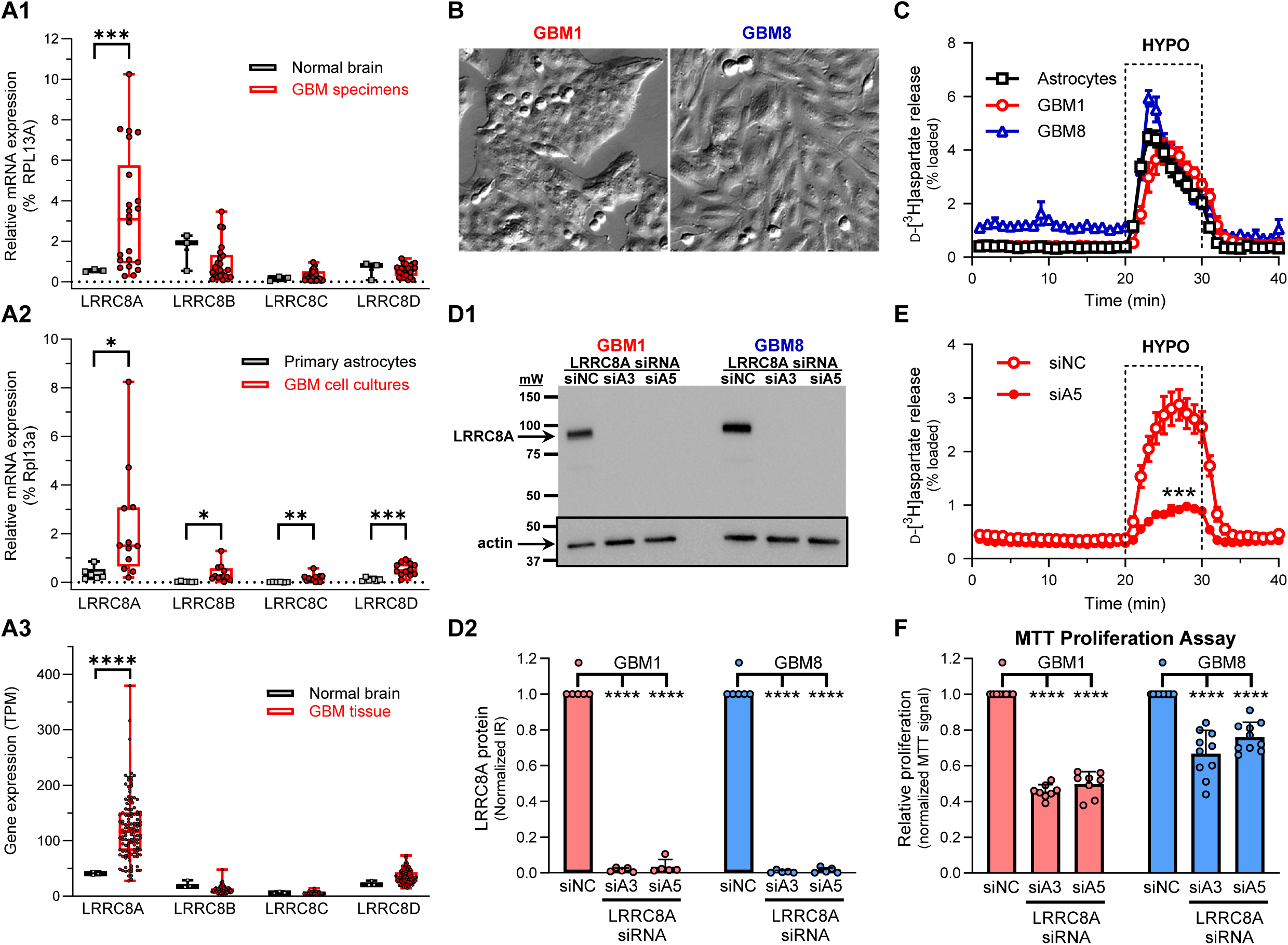
LRRC8A is upregulated in human GBM specimens and patient-derived GBM cultures and promotes cell proliferation. (**A1**) *Lrrc8* mRNA expression levels in surgically resected GBM tumor specimens (n=22) and healthy brain tissue (3 RNA medleys from 8 brains) measured using qRT-PCR and normalized to the housekeeping gene *Rpl13a*. ***p<0.001, GBM vs. normal brain. (**A2**) Comparison of *Lrrc8* mRNA expression levels in patient derived GBM cell cultures (n=12) and primary human astrocytes (n=6). *p<0.05, **p<0.01, ***p<0.001, GBM vs. astrocytes. (**A3**) In-silico analysis of RNA-seq data for the expression of *Lrrc8* isoforms in primary GBM (n=126) and healthy brain tissue (3 RNA medleys from 29 brains). GBM expression values were obtained from TCGA. Healthy brain tissue data were sourced from GEO repository, series GSE196695. ****p<0.0001, GBM vs. normal brain. In A1-A3, data are presented as boxplots with individual values and compared with an unpaired t-test with Welch correction and Holm-Šídák adjustment. (**B**) Representative images of the patient-derived GBM1 (*LEFT*) and GBM8 (*RIGHT*) cells. Images were acquired using Hoffman optics at identical 160× magnification. (**C**) VRAC activity in GBM1, GBM8, and primary human astrocytes measured as swelling activated release of preloaded D-[^3^H]aspartate. Cell swelling was induced by 30% reduction in medium osmolarity (HYPO). Mean values ±SEM from 4-6 independent experiments/cell line. (**D1**) Representative western blot image probing downregulation of LRRC8A in GBM1 and GBM8 cells treated with the LRRC8A-specific siRNAs, siA3 and siA5, and compared to the negative control siRNA (siNC). Lower inset shows the same membrane re-probed for β-actin immunoreactivity. (**D2**) Quantification of LRR8A expression in siRNA-treated GBM cultures. Mean values ±SD from 5 independent transfections/cell line. One-sample t-test with Bonferroni’s correction. ****p<0.0001 vs siNC. (**E**) Functional downregulation of VRAC activity in GBM1 cells treated with LRRC8A siRNA (siA5) and compared to siNC. Mean values ±SEM (n=5/condition). ***p<0.001 vs. siNC, t-test comparing integral D-[^3^H]aspartate release values during hypotonic exposure. (**F**) Effect of LRRC8A knockdown with the LRRC8A-specific siRNAs siA3 and siA5 on relative proliferation in GBM1 and GBM8 measured using the MTT assay. Mean values ±SD from 8-10 independent transfections. One-sample t-test with Bonferroni’s correction, ****p<0.0001 vs. siNC.

To measure VRAC activity, we used a radiotracer assay quantifying swelling-activated release of the VRAC-permeable, nonmetabolizable glutamate analogue D-[^3^H]aspartate. This method enables characterization of VRAC behavior in large cell populations and is highly suitable for cells of glial origin due to their high glutamate uptake capacity.^54,55^ Application of hypoosmotic medium, which causes cell swelling, produced a strong increase in D-[^3^H]aspartate efflux (Figure 1C) which resembled the VRAC-mediated glutamate release seen in rodent (e.g.,^54,55,56,57^) and human astrocytes (Figure 1C). The ranking of maximal release rates was GBM8 > astrocytes ≈ GBM1 (Figure 1C). Despite similarities to astrocytes, the kinetics of D-[^3^H]aspartate release in GBM had several unique features. In GBM1, maximal hypoosmotic activation occurred 3-4 min later than in GBM8 or astrocytes, indicating slower rate of cellular swelling or difference in VRAC activation kinetics (Figure 1C). In GBM8, basal (non-stimulated) release rate was approximately 3-fold higher than in GBM1 or astrocytes, likely pointing to elevated tonic VRAC activity in non-swollen GBM8 cells (Figure 1C). These differences aside, all GBM cells demonstrated functional VRAC activity.

To confirm that VRAC activity in GBM is reliant on LRRC8A, we utilized an RNAi approach. We first probed the effectiveness of four different LRRC8A-specific siRNA constructs using qRT-PCR (Figure S2A). Two effective constructs (dubbed “siA3” and “siA5”) were further validated at the protein level using Western blotting. By day 4 post-transfection, both siA3 and siA5 reduced LRRC8A protein content by >95% in both GBM1 and GBM8 (Figure 1D). These siRNAs were then tested in VRAC activity assays. In GBM1, as compared to the negative control siNC, siA3 and siA5 reduced swelling-activated D-[^3^H]aspartate release via VRAC by ≥80% (Figure 1E, Figure S2C). Strong inhibition of VRAC activity by LRRC8A siRNA was also observed in GBM8 (Figure S2D). Altogether, these results suggest that LRRC8A protein is essential for VRAC function in GBM and can be effectively downregulated by employing RNAi.

Using RNAi, we further explored the impact of LRRC8A knockdown on GBM proliferation measured as cumulative changes in cell numbers over 96 h after siRNA transfection. In MTT assays, the LRRC8A siRNAs, siA3 and siA5, reduced cell numbers by 50-55% in GBM1 and 25- 35% in GBM8 (Figure 1F). MTT results were independently confirmed using automated counting of DAPI-stained GBM nuclei. The LRRC8A-specific siA3 and siA5 strongly reduced the numbers of substrate-attached GBM1 and GBM8 cells (Figure S3). In our prior work^50^, we additionally cross-validated the effects of LRRC8A knockdown on proliferation by physically counting GBM cells using a Coulter counter. Potential contributions of cell death to the differences in cell numbers were ruled out because we found no significant increase in lactate dehydrogenase release from cells treated with siA3 or siA5 (Figure S4). As a positive control, treatment with 300 nM staurosporine caused ∼50% cell death within 24 h (Figure S4). Altogether, these results strongly support the notion that elevated LRRC8A expression in GBM boosts high rates of cell proliferation.

### The relationship between LRRC8A expression and clinical GBM outcomes

High expression of LRRC8A has been putatively linked to shorter life expectancy in several cancers (e.g.,^46,47,48,49^). Therefore, we analyzed if tumoral *Lrrc8a* mRNA levels in primary GBM correlate with clinical outcomes by performing survival analysis in TCGA datasets. The LRRC8A expression data presented in Figure 1A3 were additionally normalized using the DEseq2 variance-stabilizing transformation.^58^ This type of normalization is considered superior because it accounts for sample-to-sample differences in library size, sequencing depth, false positives, etc.^59,60^ 126 GBM TCGA patients were divided into quartiles based on their tumor *Lrrc8a* mRNA expression (Figure 2A) and analyzed for differences in their life expectancy (Figure 2B). We found that patients with the lowest *Lrrc8a* expression (Q1) had a median survival time of 480 days, which was 120+ days longer than the median survival in Q2 (333 days), Q3 (360 days), and Q4 (360 days) (Figure 2B), denoting a 33% increase in patient life expectancy. Considering the close clustering of expression levels in Q2 and Q3, we statistically compared only the highest (Q4) and the lowest (Q1) LRRC8A expression quartiles and found a trend for longer survival in Q1 (Gehan-Breslow-Wilcoxon test, p=0.078). Superficially, these results are consistent with the proliferation-promoting role for LRRC8A. However, the overall correlation between LRRC8A expression levels and survival appears to be weak, and its significance remains unclear.

**Figure 2.**
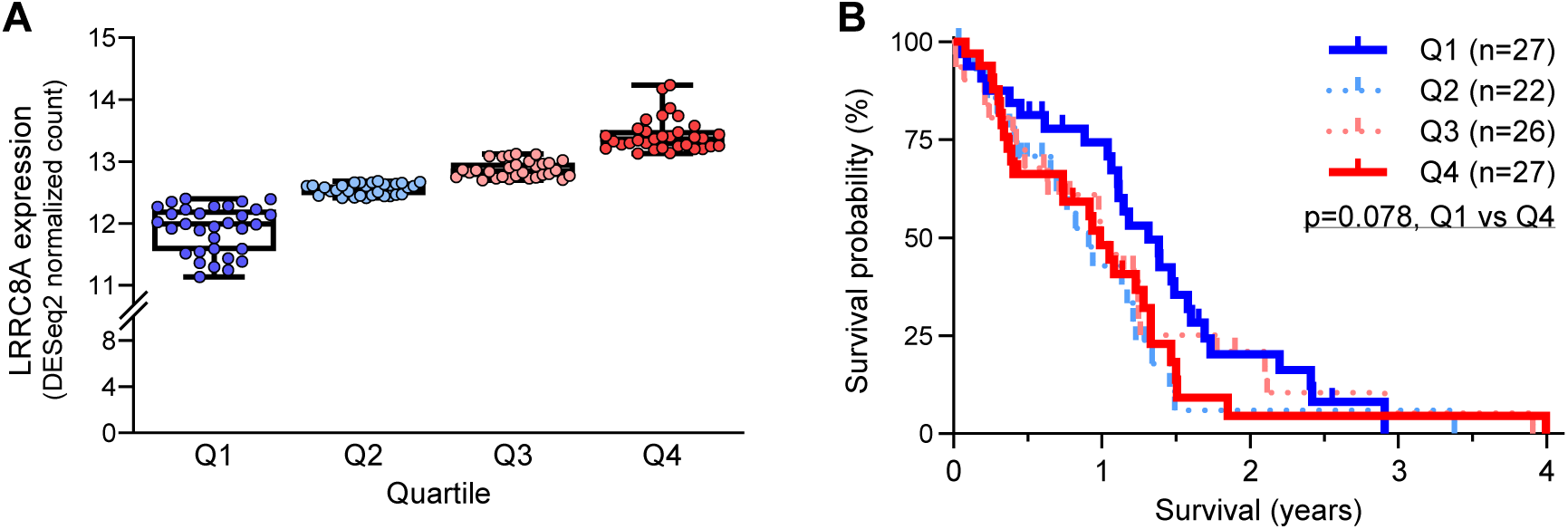
In-silico analysis of the relationship between *Lrrc8a* mRNA expression and life expectancy in GBM patients. (**A**) In-silico analysis of RNA-seq expression for *Lrrc8a* in primary GBM tissues (n=126) separated into quartiles. Data were sourced from TCGA and normalized using the DEseq2 variance-stabilizing transformation. Expression values presented as boxplots with individual values. (**B**) Kaplan-Meier plot of survival rates for GBM patients grouped by tumor *Lrrc8a* expression quartile. Median survival was compared between the lowest (Q1; n=27) and highest (Q4; n=27) *Lrrc8a* expressers using the Gehan-Breslow-Wilcoxon test. p=0.078.

### Lack of consistent effect of LRRC8A knockdown on cell cycle or cell-cycle dependent proteins

Although LRRC8A expression has been linked to high proliferative rates in several cancers (see^50,46,47,48,49^), the underlying mechanisms are far from clear and may be cancer specific. To further explore the role of LRRC8A in GBM proliferation, we started with an analysis of cell cycle progression using fluorescence-activated cell sorting (FACS) of propidium iodide-stained cells. We found significant differences between GBM1 and GBM8 cells in terms of their allocation among various cell cycle phases. In non-synchronized cultures of GBM1, ∼80%, ∼3%, and ∼10% of cells were found in G_1_/G_0_, S, and G_2_/M stages of mitotic cycle, respectively (Figure S5A,C, the remaining cells were polyploid). GBM8 demonstrated a different cell cycle distribution pattern with ∼60%, ∼4%, and ∼23% cells in G_1_/G_0_, S, and G_2_/M mitosis phases, respectively (Figure S5, the remaining cells were polyploid). Using this approach, we found no statistical differences in the apparent mitotic progression of siNC-, siA3-, or siA5-treated cells, except for a small reduction in S phase numbers in GBM8 (Figure S5B,C). The observed lack of effect of LRRC8A knockdowns on cell cycle distribution was unexpected because of the significant reduction in cell numbers seen in LRRC8A siRNA-treated cells (compare Figure S5 to the results in Figure 1F).

As a complementary approach, we evaluated the expression of cell cycle-dependent proteins using Western blotting. In specific, we analyzed expression levels for cyclin D1 and its partner CDK4, and cyclin E1 and its partner CDK2. Although we found several effects of siRNA on individual cell cycle proteins (Figure S6), we identified no common pattern of effects of LRRC8A knockdown across two different siRNA species (siA3 and siA5) and two GBM cell lines (GBM1 and GBM8). Overall, the two performed cell cycle analyses provided no clear mechanistic insight into the reduction of cell proliferation seen following LRRC8A knockdown.

### LRRC8A knockdown does not modify GRB2-dependent growth signaling

As a potential alternative mechanism underlying the observed changes in cell proliferation, we explored the effects of LRRC8A knockdown on growth signaling pathways. Previous work identified a physical interaction between LRRC8A and the growth factor signaling adaptor proteins GRB2 and GAB2.^39,42,61^ Furthermore, ablation of LRRC8A expression has been shown to reduce enzymatic activity in a diverse range of GRB2-dependent signaling cascades.^39,42,41^ Here, we focused our attention on the ERK1/2 and the JNK signaling axes, which are activated by numerous growth factor receptors and synergistically modulate AP-1 dependent gene transcription (see^62^ and diagrams in Figure 3A, F). In the case of ERK1/2 signaling, GRB2 interacts with the guanine exchange factor SOS to initiate the classical Ras➔Raf➔MEK1/2➔ Erk1/2 signal transduction (^62,63^ and Figure 3A). In the case of JNK signaling, the activation may involve the GRB2-driven Ras-Rac interactions^64^, or the PI3K-dependendent activation of a distinct SOS adaptor complex^65^ (Figure 3F). Additionally, we explored changes in the activity of the PI3K-AKT-mTOR signaling arm, which is also activated by growth factors in a GRB2/GAB2- dependent manner and is highly relevant to malignant cell proliferation, including in GBM (see ^66,67^ and diagram in Figure 4A).

**Figure 3.**
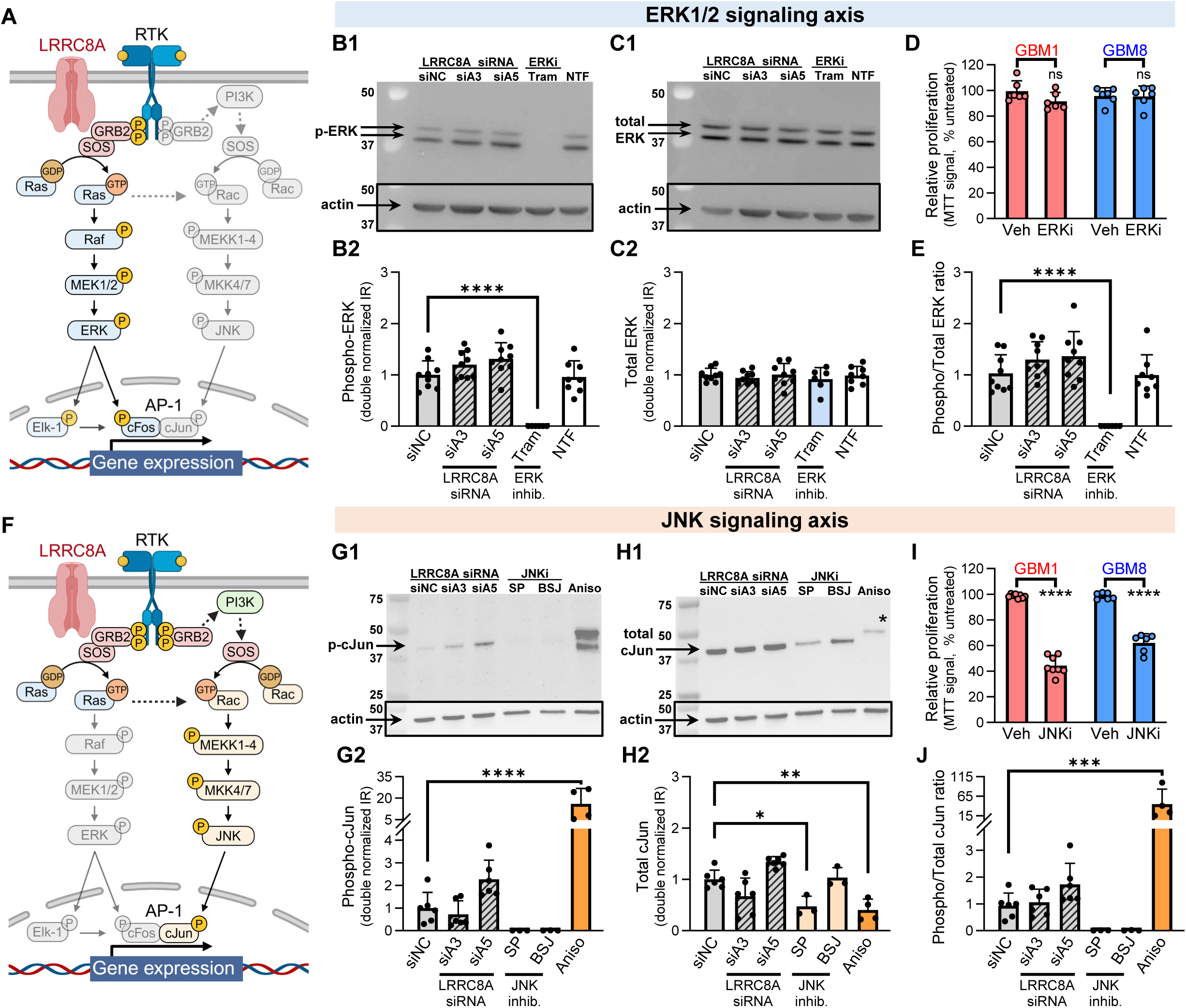
Analysis of functional interactions between LRRC8A and GRB2-dependent growth factor signaling pathways in GBM. (**A**) Diagram depicting receptor tyrosine kinase (RTK)-ERK1/2 signaling pathway and its putative interactions with LRRC8A. For abbreviations and further information see text. (**B,C,E**) Western blot analysis of the effect of LRRC8A downregulation on ERK1/2 signaling in GBM1 cells. Representative images and quantification of phospho-ERK1/2 (**B1-B2**), total ERK1/2 (**C1-C2**), and phospho/total ERK1/2 immunoreactivity ratio (**E**). GBM1 cells were treated with the LRRC8A-specific siRNAs, siA3 and siA5, and compared to the negative control siNC (n=9). GBM1 cells treated with the ERK1/2 inhibitor Trametinib (Tram; 0.3 µM, 6-hr) and non-transfected cells (NTF) were included as additional controls (n=6-8). Data are the mean values ± SD. One-way ANOVA with Dunnett’s correction, ****p<0.0001, trametinib vs siNC. (**D**) Effect of the ERK1/2 signaling inhibitor U-0126 (ERKi; 10μM) on GBM1 and GBM8 proliferation as compared to vehicle control (Veh; 0.1% DMSO). Cell proliferation values were measured using the MTT assay and normalized to within-plate untreated cells (n=6/cell line). Data are the mean values ± SD. ns, not significant, unpaired t-test. (**F**) Diagram depicting RTK-JNK signaling pathway and its putative interactions with LRRC8A. For abbreviations and further information see text. (**G,H,J**) Western blot analysis of the effect of LRRC8A downregulation on JNK signaling in GBM1 cells using cJun phosphorylation as a readout. Representative images and quantification of phospho-cJun (**G1-G2**), total cJun (**H1-H2**), and phospho/total cJun immunoreactivity ratio (**J**) in GBM1 cells treated with the LRRC8A-specific siA3 and siA5 as compared to siNC. As controls, GBM1 were treated with the JNK inhibitor SP600125 (SP, 20 µM, 24-hr), the MKK4/7 inhibitor BSJ-04-122 (BSJ, 5µM, 24-hr), or the JNK activator anisomycin (Aniso; 10µM, 2-hr). Data are the mean values ± SD from 6 siRNA experiments and 3-4 pharmacological controls. One-way ANOVA with Dunnett’s correction, *p<0.05, **p<0.01, ***p<0.001, ****p<0.0001 vs. siNC. (**I**) Effect of the JNK signaling inhibitor SP600125 (JNKi, 20µM) on GBM1 and GBM8 proliferation. The MTT assay values were normalized to within-plate untreated cells and compared to the vehicle control (Veh; 0.1% DMSO). Data are the mean values ± SD (n=6-8/cell line). Unpaired t-test, ****p<0.0001 vs. Veh.

**Figure 4.**
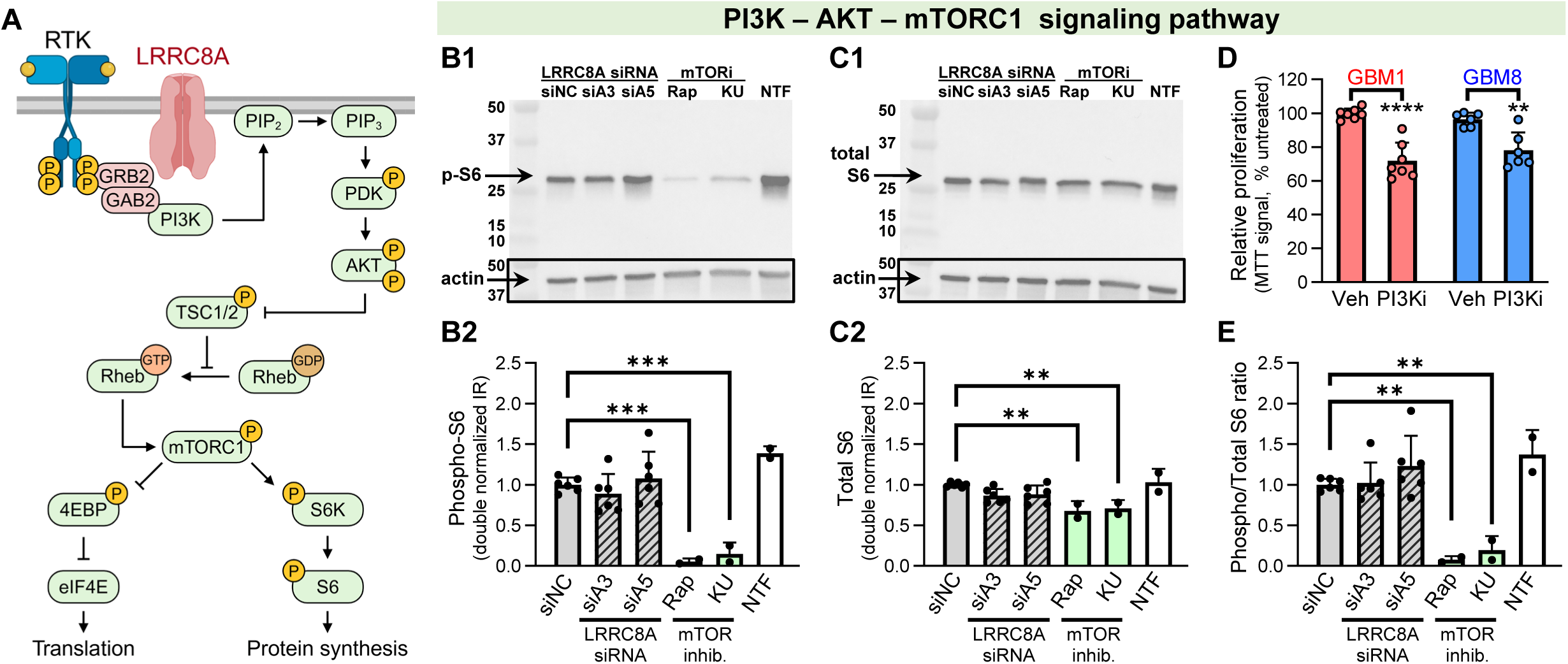
Analysis of functional interactions between LRRC8A and the GRB2-dependent PI3K/AKT/mTORC1 signaling axis in GBM. (**A**) Diagram depicting RTK-PI3K/AKT/mTORC1 signaling axis and its putative interactions with LRRC8A. For abbreviations and further information see text. (**B,C,E**) Western blot analysis of the effect of LRRC8A downregulation on mTORC1 signaling in GBM, measuring S6 phosphorylation as a readout. Representative images and quantification of phospho-S6 (**B1-B2**), total S6 (**C1-C2**), and phospho/total S6 immunoreactivity ratio (**E**). GBM1 cells were treated with the LRRC8A-specific siRNAs, siA3 and siA5, or the negative control siNC. As controls, cells were treated with the mTORC1 inhibitor Rapamycin (Rap, 10nM, 48-hr), or the mTORC1/2 blocker KU-0063794 (KU, 2.5 µM, 48-hr). NTF, non-transfected cells. Data are the mean values ± SD from 6 siRNA transfections or 2-3 pharmacological controls. One-way ANOVA with Dunnett’s correction, **p<0.01, ***p<0.001 vs. siNC. (**E**) Effect of the PI3K signaling inhibitor PI 828 (PI3Ki, 2.5μM) on GBM1 and GBM8 proliferation. The MTT assay values were normalized to within-plate untreated cells and compared to the vehicle control (Veh; 0.1% DMSO). Data are the mean values ± SD (n=6-7/cell line). Unpaired t-test, **p<0.01, ****p<0.0001 vs. Veh.

In GBM1, treatment with the LRRC8A-specific siA3 and siA5 produced no effect on phospho-Erk1/2 levels (p-Thr^202^/Tyr^204^, Figure 3B), total Erk1/2 immunoreactivity (Figure 3C), or phospho/ total Erk1/2 immunoreactivity ratio (Figure 3D). As controls, we used two inhibitors of upstream MEK1/2 kinases, U0126^68^ and trametinib^69^, both of which dramatically reduced Erk1/2 phosphorylation (Figure 3B,C,E). Surprisingly, U0126 (Figure 3D) had no significant effect on the proliferation of either GBM1 or GBM8, while trametinib produced marginal inhibition (Figure S7, 12% decrease, p<0.001). Together, these results rule out Erk1/2 signaling as the major mechanism contributing to the effect of LRRC8A on GBM proliferation.

In the next set of experiments, we analyzed the AP-1 linked JNK signaling axis by probing phosphorylation of the canonical JNK target cJun. LRRC8A knockdowns yielded no consistent effect on phospho-cJun (p-Ser^73^, Figure 3G), total cJun (Figure 3H), or phospho/total cJun immunoreactivity ratio (Figure 3J). As controls, we utilized the pharmacological inhibitors SP600125^70^ and BSJ-04-122^71^, which block JNK or the upstream kinases MKK4/7, respectively. To activate JNK, we treated cells with anisomycin. All pharmacological controls eliminated or upregulated cJun phosphorylation in the predicted manner (Figure 3G,H,J). To test the relevance of JNK signaling to GBM growth, we explored the effect of the JNK inhibitor SP600125 on GBM proliferation. Consistent with prior reports^72,73^, inhibition of JNK strongly reduced GBM cell proliferation, by 55% and 38% in GBM1 and GBM8, respectively (p<0.001, Figure 3I). This effect was quantitatively similar to the effects of LRRC8A siRNA. However, since LRRC8A knockdown caused no reduction in cJun phosphorylation, JNK signaling cannot be responsible for the pro-proliferative actions of LRRC8A.

To test for the involvement of the PI3K-AKT-mTOR pathway, we measured phosphorylation levels of one of the terminal substrates of mTORC1/S6K signaling, the ribosomal protein S6.^74^ LRRC8A knockdown yielded no effect on the levels of phospho-S6 (p-Ser^240/244^, Figure 4B), total S6 (Figure 4C), or the ratio of phospho-to-total S6 immunoreactivity (Figure 4E). As a control, we used the mTOR inhibitor rapamycin^75^, which at low nanomolar concentrations preferentially inhibits mTORC1, and the dual mTORC1/mTORC2 blocker KU-0063794^76^. Both agents effectively reduced S6 phosphorylation by 85-95% (Figure 4B,E) confirming the specificity and the sensitivity of this assay. Finally, we tested the effect of the PI3K inhibitor PI 828^77^ on GBM proliferation. This agent reduced GBM1 and GBM8 cell numbers by 28% and 19%, respectively (Figure 4D). In summary, the lack of effect of LRRC8A siRNA on phospho-S6 levels rules out the PI3K-dependent mTORC1 signaling as the basis for pro-proliferative actions of LRRC8A in GBM.

### LRRC8A knockdown reduces [Cl^−^]_i_ and modifies the Cl^−^-sensitive WNK-SPAK/OSR1 axis

As the next alternative, we explored if GBM proliferation is regulated by VRAC-mediated Cl^−^ fluxes and/or changes in intracellular Cl^−^ levels ([Cl^−^]_i_). The activity of several Cl^−^ channels and the associated changes in [Cl^−^]_i_ have been linked to progression through the mitotic cycle and cell migration, including in GBM cell lines.^78,79^ We first assessed [Cl^−^]_i_ in GBM by non-invasively measuring steady-state distribution of the radiotracer ^36^Cl^−^ (see *Star*⋆*Methods* and ref.^80,81^ for implementation in various cell types). In GBM1, ^36^Cl^−^ uptake reached equilibrium within ∼10 min (Figure 5A). Therefore, all subsequent experiments determined steady-state ^36^Cl^−^ content after 20 min of incubation with this radiotracer. Using this approach, we found that LRRC8A knockdown reduced [Cl^−^]_i_ by 5-8% (Figure 5B). Although small, this effect was significant and consistent among two different siRNA constructs (8% for siA3 and 5% for siA5, p<0.0001 and p<0.01, respectively, Figure 5B). Thus, activity of the LRRC8A-containing VRACs appear to sustain high [Cl^−^]_i_. To further explore if VRAC activity modifies GBM proliferation rate, we tested the putative VRAC blocker DIDS.^82,83^ The advantage of DIDS is that it is non-toxic in longitudinal cell proliferation experiments, unlike DCPIB^84^. Within a 72-h incubation period, DIDS reduced GBM1 proliferation by 49% (p<0.001) and GBM8 proliferation by 22% (p=0.019), respectively (Figure 5C). The lower efficacy in GBM8 is likely due to limited inhibition of VRAC (23% inhibition of channel activity, p=0.015, Figure S8). DIDS blocks VRAC poorly at negative voltages and may be less efficacious in cells with highly negative membrane potential.^82,83^ With this caveat notwithstanding, our pharmacological data support the notion that VRAC modulates GBM proliferation.

**Figure 5.**
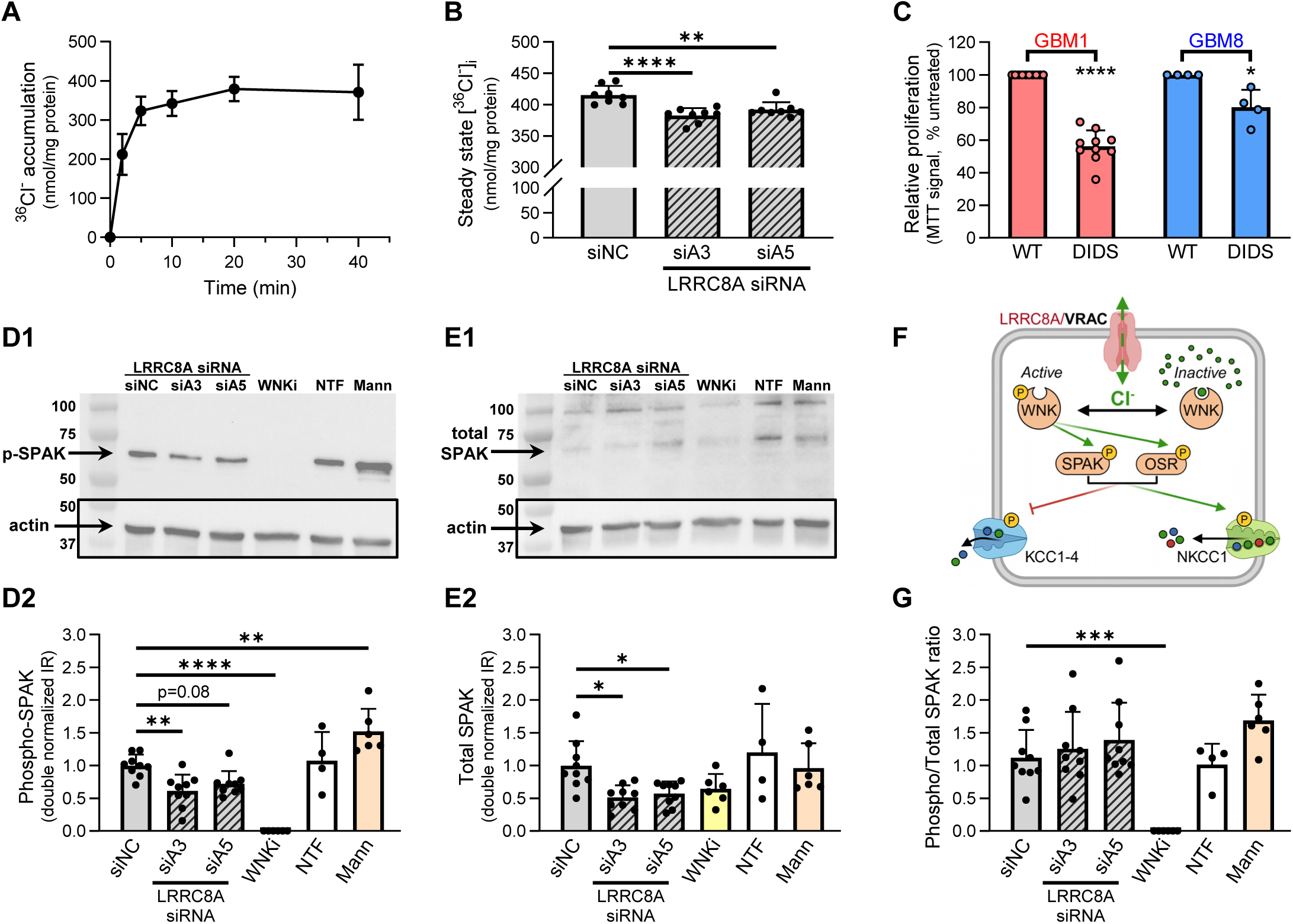
Effect of LRRC8A knockdown on [Cl^−^]_i_ and the Cl^−^-sensitive WNK signaling axis. (**A**) Kinetics of ^36^Cl^−^ accumulation in GBM1 cells measured in a serum-containing cell culture medium. Data are the mean values ± SD of 4 independent assays. (**B**) Effect of LRRC8A knockdown on a steady state ^36^Cl^−^ accumulation in GBM1 cells. Cells were transfected with the LRRC8A-specific siRNAs, siA3 and siA5, or the negative control siNC. 96-h later, intracellular Cl^−^ levels were measured as steady state ^36^Cl^−^ uptake. Data are the mean values ± SD from 8 independent transfections. One-way ANOVA with Dunnett’s correction. **p<0.01, ****p<0.0001 vs. siNC. (**C**) Effect of the putative VRAC blocker DIDS (500 µM) on GBM1 and GBM8 cell proliferation as compared to untreated cells (WT). Proliferation was quantified using an MTT assay and normalized to untreated cells on the same plate. Data are the mean ±SD. n=4-10/cell line. One-sample t-test. *p<0.05, ****p<0.0001 vs WT. (**D,E,G**) Western blot analysis of the effect of LRRC8A downregulation on WNK signaling in GBM1, measured using SPAK phosphorylation as a readout. Representative images and quantification of phospho-SPAK (**D1-D2**), total SPAK (**E1-E2**), and phospho/total SPAK immunoreactivity ratio (**G**). GBM1 were treated with the LRRC8A-specific siA3 and siA5 or the negative control siNC (n=9). As controls, cells were treated with the WNK inhibitor WNK463 (‘WNKi’, 1µM, 24-hr) or hyperosmotic medium supplemented with 300 mM mannitol (‘Mann’, 1- hr treatment). NTF, non-transfected cells. n=4-6 for pharmacological controls. Data are the mean ±SD. One-way ANOVA with Dunnett’s correction, *p<0.05, **p<0.01, ***p<0.001, **** p<0.0001 vs. siNC. (**F**) Hypothetical diagram linking VRAC activity to the activity of the Cl^−^ sensitive WNK kinases and intracellular Cl^−^ homeostasis. The downstream protein kinases SPAK and OSR1 regulate activity of cation-chloride cotransporters NKCC1 and KCC1-4, which contribute to [Cl^−^]_i_ control.

Since [Cl^−^]_i_ is inversely related to the activity of the Cl^−^-sensitive WNK kinases and their downstream targets SPAK and OSR1^85^, we further investigated if LRRC8A knockdown modulates WNK signaling. WNK activity was determined by measuring phosphorylation levels of the downstream target SPAK (p-Ser^373^), with the caveat that the antibody used in this analysis additionally recognizes phosphorylation of the structurally and functionally homologous OSR1 (p-Ser^325^). The LRRC8A-targeting siA3 and siA5 constructs reduced levels of phosphorylated SPAK/OSR1, by 40% (p=0.002) and 26% (p=0.077) respectively (Figure 5D). The effect was mirrored by the loss of immunoreactivity for total SPAK protein (a decrease of 46% by siA3, p=0.015, and 40% by siA5, p=0.047, Figure 5E) resulting in an unchanged phospho-SPAK/total SPAK ratio (Figure 5F). The specificity of immunoblotting signals was further validated by measuring the effects of a WNK inhibitor and WNK activator. The pan-WNK inhibitor WNK463 eliminated SPAK/OSR1 phosphorylation (Figure 5D, p<0.001). Conversely, activation of WNKs by incubation in 600 mOsm hyperosmotic medium^86^ increased phospho-SPAK/OSR1 signal (+52%, p<0.001, Figure 5D) and phospho/total SPAK immunoreactivity ratio (+83%, p=0.015, Figure 5F). The intermediate conclusion from these results is that LRRC8A knockdown reduces total SPAK (and likely OSR1) levels without impacting relative WNK-SPAK/OSR1 activity.

### The non-enzymatic contributions of WNK1 to mTORC2 signaling and GBM proliferation

Since LRRC8A knockdown reduced SPAK expression, we further investigated if modified WNK-SPAK/OSR1 signaling might underlie the observed changes in GBM proliferation. Such an idea is consistent with prior literature reports on the important role of WNKs, including WNK1, in cell proliferation and numerous aspects of cancerogenesis (reviewed in^87,88^). We first checked if pharmacological inhibition of WNK suppresses GBM growth. The pan-inhibitor of WNK isozymes, WNK463^89^, eliminated phosphorylation of downstream SPAK and OSR1 at 1 µM (Figure 5D,G) but was completely ineffective in reducing GBM proliferation across a broad concentration range of 0.1-3 µM (Figure 6A). Still, several alternative, non-enzymatic, non-SPAK/OSR1-dependent mechanisms have been proposed for WNK1 including regulation of mitotic division^90^ and regulation of the pro-growth SGK1 signaling^91,92^. Therefore, we tested the effects of WNK1 knockdown using three different siRNA constructs (Figure 6B). The WNK1- targeting siRNAs reduced proliferation in both GBM1 and GBM8, in a manner that was commensurate with their effect on WNK1 mRNA expression (Figure 6C, compare to siRNA potencies in Figure 6B). The most effective WNK1 siRNA construct, termed #5, inhibited GBM1 and GBM8 proliferation by 70% and 47%, respectively (p<0.001, Figure 6C) exceeding the anti-proliferative effects of LRRC8A knockdown.

**Figure 6.**
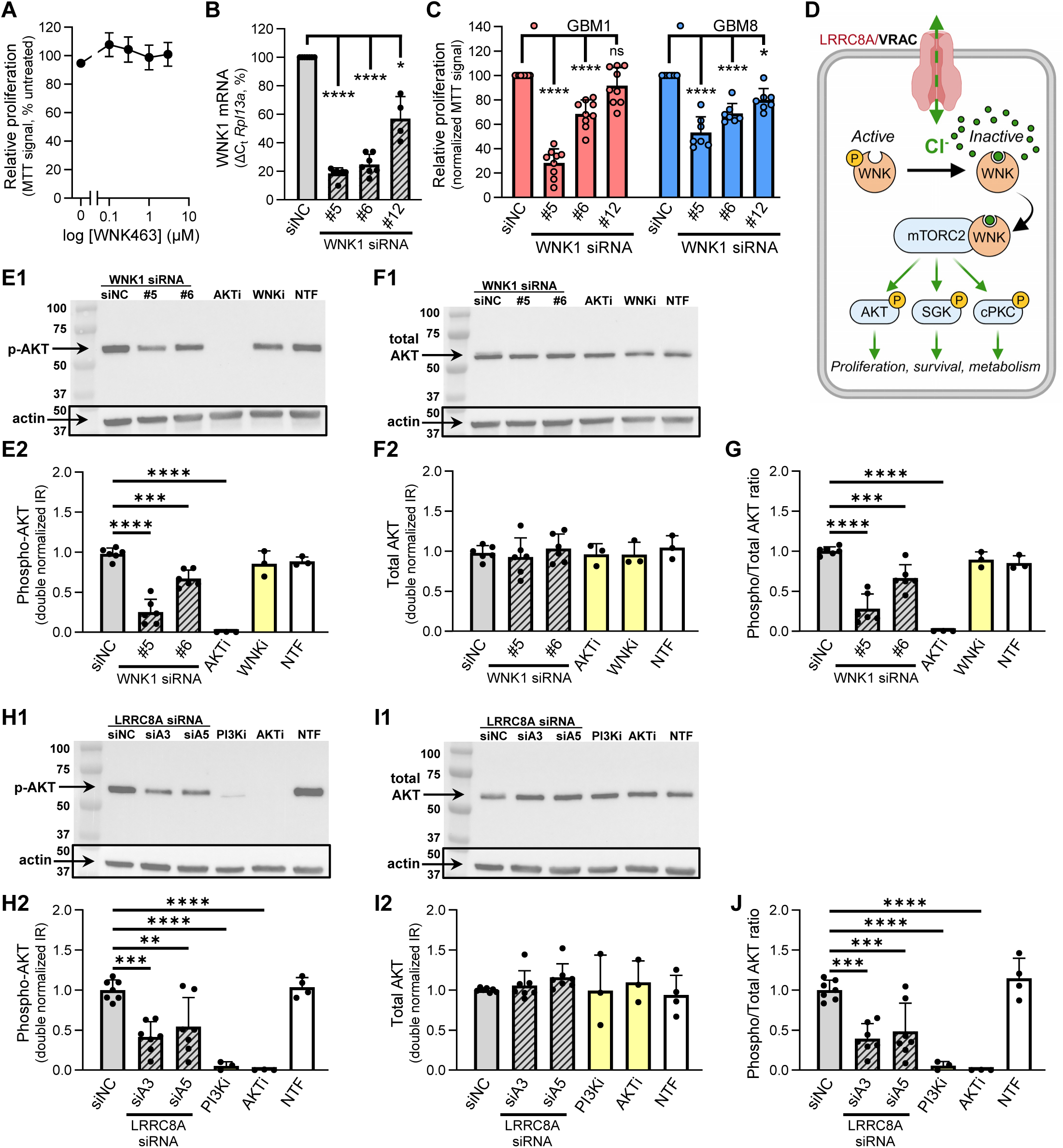
LRRC8A knockdowns regulates mTORC2 signaling through WNK1-dependent mechanism. (**A**) Effect of WNK signaling inhibitor (WNK463, 0.1-3µM) on GBM1 proliferation. Proliferation rates were measured using the MTT assay and compared to vehicle control (0.1% DMSO). Data are the normalized mean values ± SD from 3 independent treatments. (**B**) Effect of WNK1 knockdown on *Wnk1* mRNA levels. GBM1 cells were transfected with WNK1-specific siRNAs #5, #6, and #12 or the negative control siNC. Expression values were measured using qRT-PCR and normalized to *Rpl13a*. Data are the mean values ± SD. n=7 independent transfections. One-sample t-test with Bonferroni correction, *p<0.05, ****p<0.0001, vs. siNC. (**C**) Effect of WNK1 knockdown on proliferation in GBM1 and GBM8. Cells were treated with WNK1-specific siRNAs (#5, #6, and #12) or the negative control siNC. Proliferation was measured using an MTT assay. Data are the normalized mean values ±SD from 7-9 independent transfections/cell line. One-sample t-test with Bonferroni correction, ****p<0.0001 vs. siNC. (**D**) Diagram depicting hypothetical interaction between LRRC8A/VRAC, the Cl^−^-dependent WNK1, and mTORC2 signaling in GBM. VRAC promotes higher [Cl^−^]_i_ and formation of the signaling complex between the Cl^−^-bound WNK1 and mTORC2. mTORC2 activation promotes GBM proliferation via the downstream protein kinases AKT, SGK, and/or PKC. (**E-G**) Western blot analysis of the effect of WNK1 downregulation on mTORC2 signaling in GBM1 cells, measured using AKT phosphorylation as a readout. Representative images and quantification of phospho-AKT (**E1-E2**), total AKT (**F1-F2**), and phospho/total AKT immunoreactivity ratio (**G**). GBM1 cells were transfected with WNK1-specific siRNAs (#5, #6, and #12) or the negative control siNC (n=6). As additional controls (n=3), GBM1 were treated with the AKT inhibitor MK-2206 (AKTi, 2.5 µM, 24 h) and the WNK inhibitor WNK463 (WNKi, 1µM, 24 h). NTF, non-transfected cells. Data are the mean values ±SD. One-way ANOVA with Dunnett’s correction, ***p<0.001, **** p<0.0001 vs. siNC. (**H-J**) Western blot analysis of the effect of LRRC8A downregulation on mTORC2 signaling in GBM, using AKT phosphorylation as a readout. Representative images and quantification of phospho-AKT (**H1-H2**), total AKT (**I1-I2**), and phospho/total AKT immunoreactivity ratio (**J**). GBM1 were treated with siA3, siA5, or siNC (n=7). Non-transfected cells (NTF) and cells treated with the PI3K inhibitor PI 828 (PI3Ki, 2.5 µM, 24 h) or the AKT inhibitor MK-2206 (AKTi, 2.5 µM, 24 hr) were included as additional controls (n=3-4). Data are the mean values ± SD. One-way ANOVA with Dunnett’s correction, **p<0.01, ***p<0.001, ****p<0.0001 vs. siNC.

As the potent WNK inhibitor WNK463 did not affect cell proliferation (see Figure 6A), we probed for the potential involvement of a newly discovered mechanism which involves the formation of a complex between the enzymatically inactive, Cl^−^-bound WNK1 and mTOR complex 2 (mTORC2).^92^ mTORC2 activity was assessed by measuring the phosphorylation levels of one of its substrates, AKT (phospho-Ser^473^).^93^ Two efficacious WNK1 siRNAs, construct #5 and construct #6, decreased the mTORC2-dependent phosphorylation of AKT by 74% (p<0.001) and 31% (p<0.001), respectively, in a manner that was proportional to their effect on WNK1 mRNA levels (compare to Figure 6B). There was no effect on total AKT levels (Figure 6F). Accordingly, changes in the phospho/total AKT immunoreactivity ratio (Figure 6G) mirrored the pattern of changes in phospho-AKT. As a positive control, we used the AKT inhibitor MK-2206^94^, which abolished AKT phosphorylation (p<0.001, Figure 6E, G).

To test if the same signaling changes occurred following LRRC8A knockdown, we quantified levels of AKT phosphorylation in LRRC8A siRNA-treated GBM1 cells. The LRRC8A-targeting siA3 and siA5 constructs both reduced mTORC2-dependent phospho-AKT immunoreactivity, by 59% (p<0.001) and 45% (p=0.003) respectively (Figure 6H, J). There were no changes in total AKT expression levels (Figure 6I). As controls, we employed the PI3K inhibitor PI 828 and the AKT inhibitor MK-2206. Both agents eliminated AKT phosphorylation (95+% reduction in phospho-signal) without affecting total AKT levels (Figure 6H-I). Together, these results strongly suggest that LRRC8A protein expression and/or VRAC function strongly modulate the activity of mTORC2.

### mTORC2 signaling drives GBM proliferation

To test the plausibility of the notion that decreased mTORC2 signaling is what causes impaired proliferation in LRRC8A knockdown cells, we measured the impact of mTORC2 signaling on GBM proliferation, first by blocking mTOR activity with the mTORC1/2 inhibitor, KU-0063794.^76^ This compound potently reduced proliferation of GBM1 and GBM8 by 58% (p<0.001) and 45% (p<0.001), respectively (Figure 7A, B). We next tested the effects of pharmacological inhibitors targeting the three canonical substrates of mTORC2 catalytic activity: AKT (MK-2206), SGK (GSK650394)^95^, and conventional PKCα/β/γ (cPKC, Gö6976). These AGC family protein kinases are known to promote proliferation in numerous cell types (e.g.,^96^). When used individually, the AKT and SGK blockers inhibited cell growth in the range of 10-30% in both patient-derived cell lines (Figure 7A, B, p<0.001). The cPKC inhibitor was completely ineffective (Figure 7A, B). However, when all three agents were combined, proliferation was reduced by 53% and 42% in GBM1 and GBM8, respectively (Figure 7A, B, p<0.001). The apparent additive action of AKT and SGK inhibitors closely matched the effects of the mTOR blocker KU-0063794 in both GBM cell lines (Figure 7A, B). Collectively, the results of our siRNA and pharmacological experiments support the hypothesis that LRRC8A regulates GBM proliferation through activation of the mTORC2 signaling axis.

**Figure 7.**
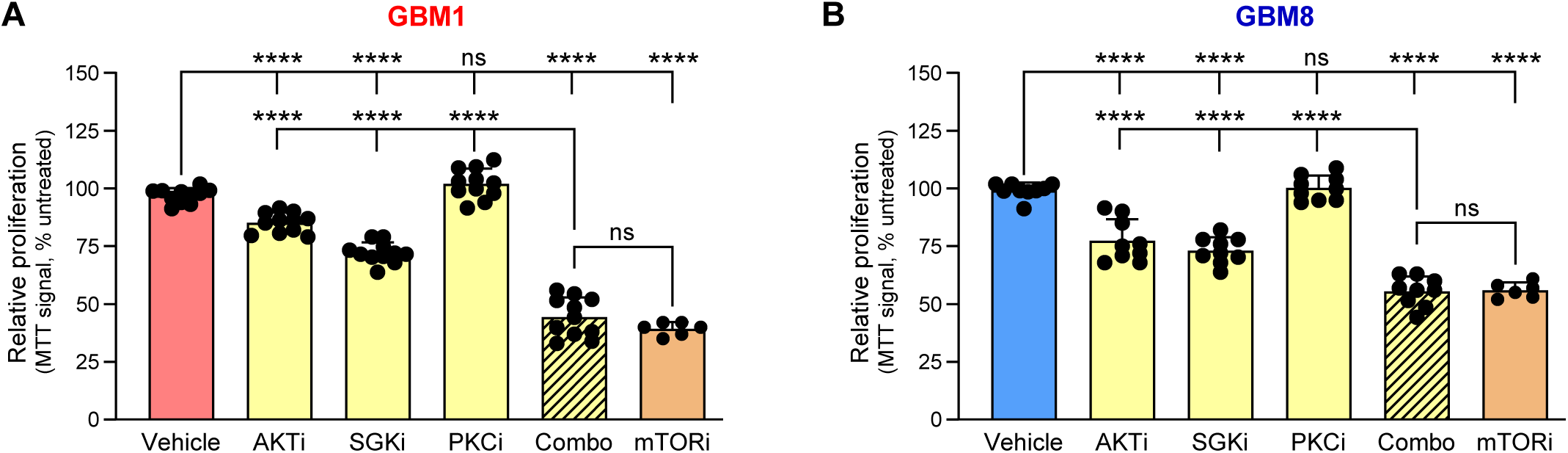
Effect of mTORC2-dependent signaling inhibition on GBM cell proliferation. (**A**) GBM1 cells were treated for 3 days with the inhibitors of AKT (AKTi; MK-2206, 2.5 μM), SGK (SGKi; GSK650394, 10 μM), PKC (PKCi; Gö6976, 3 μM), a combination of AKTi/SGKi/PKCi (Combo), or mTORC1/2 (mTORi; KU-0063794, 2.5 μM). The resulting effect on cell proliferation was measured using an MTT assay and compared to vehicle-treated cells (Veh; 0.167% DMSO). Data are the mean values ± SD from 6-9 independent treatments. One-way ANOVA with Bonferroni correction. ns, not significant, ****p<0.0001. (**B**) The same pharmacological treatments as shown in panel A were evaluated for their effects on proliferation of GBM8 cells (n=6-9). One-way ANOVA with Bonferroni correction. ns, not significant, ****p<0.0001.

## DISCUSSION

Three major findings of this study can be summarized as follows: (i) The Cl⁻ channel-forming protein LRRC8A shows a pattern of strongly elevated expression in human GBMs. (ii) RNAi analysis in patient-derived GBM cultures indicates that LRRC8A facilitates cell proliferation. (iii) the influence of LRRC8A on proliferation is mediated by a novel Cl⁻-sensitive signaling mechanism involving a non-enzymatic function of WNK1 and its impact on downstream mTORC2 signaling. We believe that the results reported here will be of significant interest to neuro-oncologists, as well as to the broader community of cell physiologists exploring the diverse functions of volume-regulated anion channels and the VRAC-constituting LRRC8 proteins.

This study is the first to report that LRRC8A is overexpressed in the majority of primary GBMs as compared to healthy brain tissue. The pattern of strongly elevated LRRC8A expression was consistently identified in surgical GBM specimens, patient-derived GBM cultures, and in the GBM datasets deposited in TCGA. These findings may suggest a pathological role for LRRC8A in GBM progression, with potential implications for clinical outcomes. Similar to our current work, elevated LRRC8A expression, compared to adjacent non-cancerous tissues, has been found in colon cancer, esophageal squamous cell carcinoma, hepatocellular carcinoma, gastric cancer, and pancreatic adenocarcinoma.^46,47,48,97,98^ In all of these carcinomas, as well as in head-and-neck cancer, elevated LRRC8A expression was predictive of poor survival outcomes, suggesting that it may serve as a biomarker for aggressive tumors.^46,47,48,97,98,99^ In contrast, in a systematic analysis of TCGA datasets, Carpanese et al. identified the opposite trend in kidney renal clear cell carcinoma, low-grade gliomas, and sarcomas, where high LRRC8A levels were associated with better survival outcomes.^99^ In GBM, we found a weak correlation between LRRC8A expression and patients’ life expectancy, with a trend for longer survival in the patient cohort with the lowest LRRC8A levels. Nevertheless, our data suggest that LRRC8A protein is necessary for the effective proliferation of GBM cells. This reliance on LRRC8A was confirmed in patient-derived GBM cells expressing either low (GBM1) or high (GBM8) levels of LRRC8A. We acknowledge that the dependence on LRRC8A is likely a cell type-specific phenomenon since LRRC8A deletion yielded no effect on proliferation in several non-malignant and malignant cell lines (e.g.,^100^).

Across the existing literature, no unifying mechanism for LRRC8A-dependent cancer growth has been agreed upon. Pioneer work in immune cells revealed a physical interaction between LRRC8A, GRB2, and GAB2.^39^ These two scaffolding proteins transduce growth factor and cytokine receptor signaling and are critical for normal development, as well as for the proliferation and motility of cancer cells.^101,102,66^ Deletion or knockdown of LRRC8A has been shown to reduce enzymatic activity in the growth factor-dependent ERK1/2 and PI3K-AKT signaling pathways in thymocytes, myocytes, endothelial cells, and other diverse cell types.^39,41,42^ In our hands, LRRC8A knockdown did not cause changes in the activity of three GRB2-dependent signaling arms, namely ERK1/2, JNK, and PI3K-AKT-mTORC1. Consistent with our data in GBM, LRRC8A downregulation in hepatocellular carcinoma did not reduce relative activity of ERK1/2 or AKT.^47^ Thus, regulation of GRB2-dependent signaling by LRRC8A appears to be a cell type-specific phenomenon that must be tested and confirmed on a case-by-case basis. Among other suggested mechanisms, several groups have reported that LRRC8A depletion causes disruptions in cell cycle-related signaling, such as upregulation of the CDK inhibitors p21^Cip1^ and p27^Kip1^ or downregulation of cyclin D1 and CDK2, in addition to increased pro-apoptotic signaling and cell death.^46,47,48^ In our study, despite the strong reduction in GBM cell numbers, LRRC8A knockdown did not produce consistent changes in cell cycle distribution, levels of cell cycle-related proteins, or increased apoptosis. These findings are mirrored by a recently published extensive analysis of the effect of LRRC8A knockout in colorectal carcinoma HCT116 cells, where a strong reduction in cell proliferation was not associated with changes in cell cycle distribution or apoptosis.^99^ Overall, the mechanistic link between LRRC8A and cell proliferation remains obscure.

In search of an alternative explanation, we explored the impact of reduced VRAC channel activity on [Cl^−^]_i_ homeostasis. Over years, many laboratories have found that structurally diverse VRAC blockers inhibit proliferation in various cell types, including GBM (e.g.,^28,33,103,104^ and reviews^26,27,105^). However, exceptions do exist.^100^ For example, in C2C12 myoblasts neither pharmacological inhibition of VRAC or LRRC8A knockdown impair cell growth, but instead potently inhibit cell differentiation.^106^ In our hands, the putative VRAC blocker DIDS has reduced GBM1 and GBM8 proliferation with the inhibitory potency resembling that of LRRC8A knockdown (GBM1>GBM8). The similarity of our pharmacology and molecular biology results, combined with existing published data, affirm that LRRC8A contributes to GBM proliferation through its role in Cl^−^ channel function rather than as a signaling scaffold protein. Yet, interpretations of the effects of VRAC blockers have been called into question due to the limited specificity and/or toxicity of these compounds.^107^ Complementary evidence for the importance of [Cl^−^]_i_ in cell proliferation can be found in the anti-proliferative effects of low-Cl^−^ media.^48,108,109^ In the context of GBM, our functional assays showed that LRRC8A knockdown not only potently suppressed VRAC activity but also reduced [Cl^−^]_i_. A similar effect of LRRC8A downregulation on [Cl^−^]_i_ and proliferation was observed in gastric cancer cells.^48^ These findings may have mechanistical relevance because recent studies discovered molecular machinery enabling [Cl^−^]_i_- dependent modulation of cellular functions, principally through the atypical kinases from the With-No-lysine(K) family (WNK1-4).^110,111^ In WNK isozymes, particularly WNK1 and WNK4, Cl^−^ binds to the autoinhibitory C-terminal domain and prevents enzymatic activity.^112,113^ Consequently, when [Cl^−^]_i_ is reduced, WNKs undergo autophosphorylation and become active. Importantly, WNK1 has emerged as a prominent player in progression of several cancers and is thought to contribute to numerous aspects of tumorigenesis.^87,88,111^ Based on the reduction of [Cl^−^]_i_ in LRRC8A knockdown cells, we expected that WNK activity would be increased. Yet, we found no evidence indicating any such effect in our phosphoproteomics assays. Furthermore, the potent WNK inhibitor WNK463 had no influence on cell proliferation despite complete inhibition of WNK activity.

Besides its well-known protein kinase actions, WNK1 may modulate intracellular signaling via a non-enzymatic mechanism (e.g.,^91,92^). Therefore, we additionally tested for the effects of WNK1 siRNA. In striking contrast to the lack of effect of pharmacological inhibitor, WNK1 siRNA knockdown strongly suppressed GBM proliferation. The potential underlying process has been identified only recently. Saha et al.^92^ found that the Cl^−^-bound WNK1 activates mTORC2 through a previously unknown scaffolding mechanism leading to activation of downstream AGC protein kinases, such as SGK1, AKT, and conventional isoforms of PKC.^114^ We hypothesize that, in GBM cells, LRRC8A-containing VRAC sustains elevated [Cl^−^]_i_ and facilitates the Cl^−^-dependent WNK1/mTORC2 scaffolding and downstream signaling. We probed and validated this mechanism using the mTORC1/2 inhibitor KU-0063794 and, alternatively, by blocking the mTORC2-dependent AKT and SGK. Inhibition of mTOR activity, or combinatorial inhibition of AKT and SGK, fully recapitulated the effect of LRRC8A knockdown, thus firmly supporting the new signaling mechanism presented in Figure 8.

**Figure 8.**
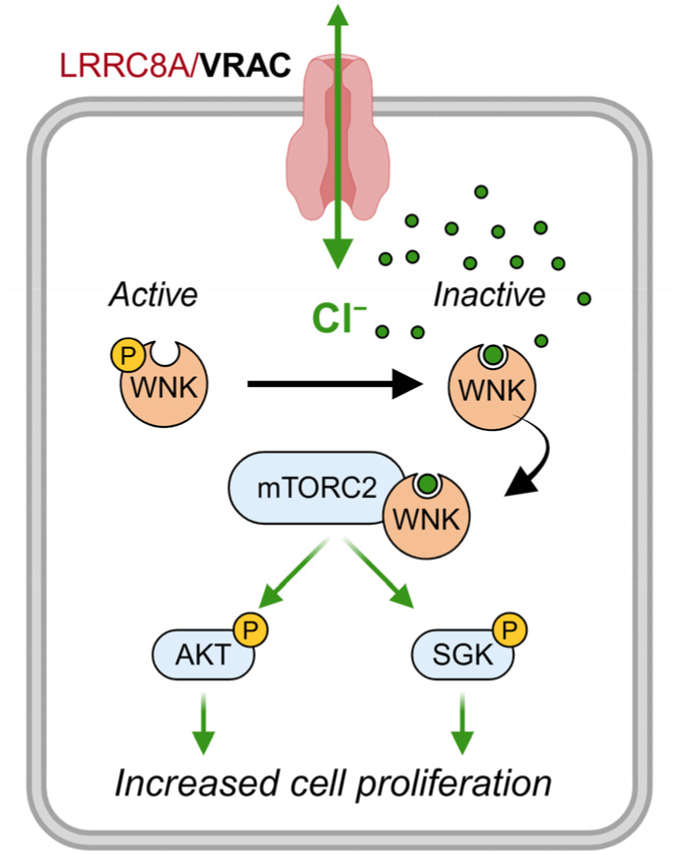
LRRC8A-containing VRAC regulates GBM proliferation through a WNK1/ mTORC2-dependent mechanism. Hypothetical model implicating the Cl^−^-dependent WNK1/mTORC2 signaling as the mechanistic link between LRRC8A expression, VRAC activity, and GBM proliferation. VRAC activity is proposed to sustain high [Cl^−^]_i_, promote formation of the complex between Cl^−^-bound WNK1 and mTORC2. The subsequent increase in mTORC2 activity promotes GBM proliferation through activation of homologous AKT and SGK.

### Limitations of the study

In this study, we identified and tested a novel mechanism linking the LRRC8A-containing anion channel VRAC to WNK1/mTORC2-dependent GBM proliferation. There are some limitations of this work, which may be further addressed in future studies. (i) It will be important to probe if the proposed signaling pathway modulates cell growth in a universal or tissue/cell type specific fashion. At this stage there are two additional precedents for the WNK1/mTORC2-dependent regulation of cellular functions in renal tubule cells^92^ and myofibroblasts^81^. (ii) It would be helpful to confirm the selective involvement of mTORC2 in the LRRC8A-dependent GBM proliferation using tools of molecular biology (e.g., RNAi) and and perform rescue experiments verifying that LRRC8A, WNK1, and mTORC2 act via a shared mechanism. (iii) One may want to additionally probe for physical interactions between WNK1 and mTORC2 in GBM (such interaction has been firmly established in renal tubule cells^92^). (iv) The transcriptional elements involved in the LRRC8A/WNK1/mTORC2-dependent regulation of GBM proliferation would have to be further explored. (v) The present findings will have to be recapitulated in animal cancer models, which better reflect the complex tumor microenvironment. Finally, (vi) our findings have been thoroughly tested in two patient-derived GBM cell lines only. Although diverse, these cell lines are unlikely to reflect the entire spectrum of GBM heterogeneity. While these caveats can be addressed in future work, they do not alter the major conclusions of the present manuscript.

## DECLARATION OF GENERATIVE AI AND AI-ASSISTED TECHNOLOGIES IN WRITING PROCESS

Authors declare that no generative AI or AI-assisted technologies were used in preparing this manuscript. Authors take full responsibility for the content of this publication.

## STAR⋆METHODS

Detailed methods are provided in the online version of this paper and include the following:

- KEY RESOURCES TABLE
- RESOURCES AVAILABILITY

- Lead contact
- Materials and availability statement
- Data and code availability
- EXPERIMENTAL MODEL AND STUDY PARTICIPANTS DETAILS

- Patient samples
- Primary GBM cell lines
- The Cancer Genome Atlas (TCGA) datasets
- METHOD DETAILS

- Computational analysis of TCGA dataset
- mRNA isolation and real-time quantitative PCR
- Protein isolation and Western blot analysis
- siRNA manipulation of gene expression
- Radioisotope assay of VRAC activity
- Cell proliferation assays
- Measurements of intracellular Cl^−^ levels
- Fluorescence-activated cell sorting (FACS) analysis
- QUANTIFICATION AND STATISTICAL ANALYSIS

- Statistical analyses

## SUPPLEMENTAL INFORMATION

The primary data analyzed in this study have been deposited in Open Science Framework and can be found online at https://osf.io/2r7k9

## ACKNOWLEDGEMENTS

We are grateful to Drs. David Jourd’heuil, Corinne S. Wilson, and Ryan Kanai for their help with pilot experiments leading to this work. This study was supported in part by the Dr. Louis Sklarow Memorial Trust, the U.S. National Institutes of Health (grants R01 NS111943 to A.A.M. and R01 HL149993 to N.O.D.), and a translational grant awarded by Albany Medical College (to Y.H.K and A.A.M.).

## AUTHOR CONTRIBUTIONS

A.A.M., A.M.F., and M.D.B. conceptualized this study. A.M.F., M.D.B., S.O., N.M., M.A.B., and A.A.M. conducted all included experiments. A.M.F., M.D.B., S.O., I.F.A., A.R.P., Y.H.K., and A.A.M. procured and generated GBM specimens and GBM cell lines. A.M.F., M.D.B., S.O., N.M., A.P.A., M.A.B., N.O.D. and A.A.M. performed data analysis and interpretation. A.A.M., N.O.D., and Y.H.K. acquired funds to support this study. A.A.M. and A.M.F. prepared the initial draft of the manuscript. All authors contributed to manuscript writing and revisions.

## STAR*METHODS (full online version)

### KEY RESOURCES TABLE

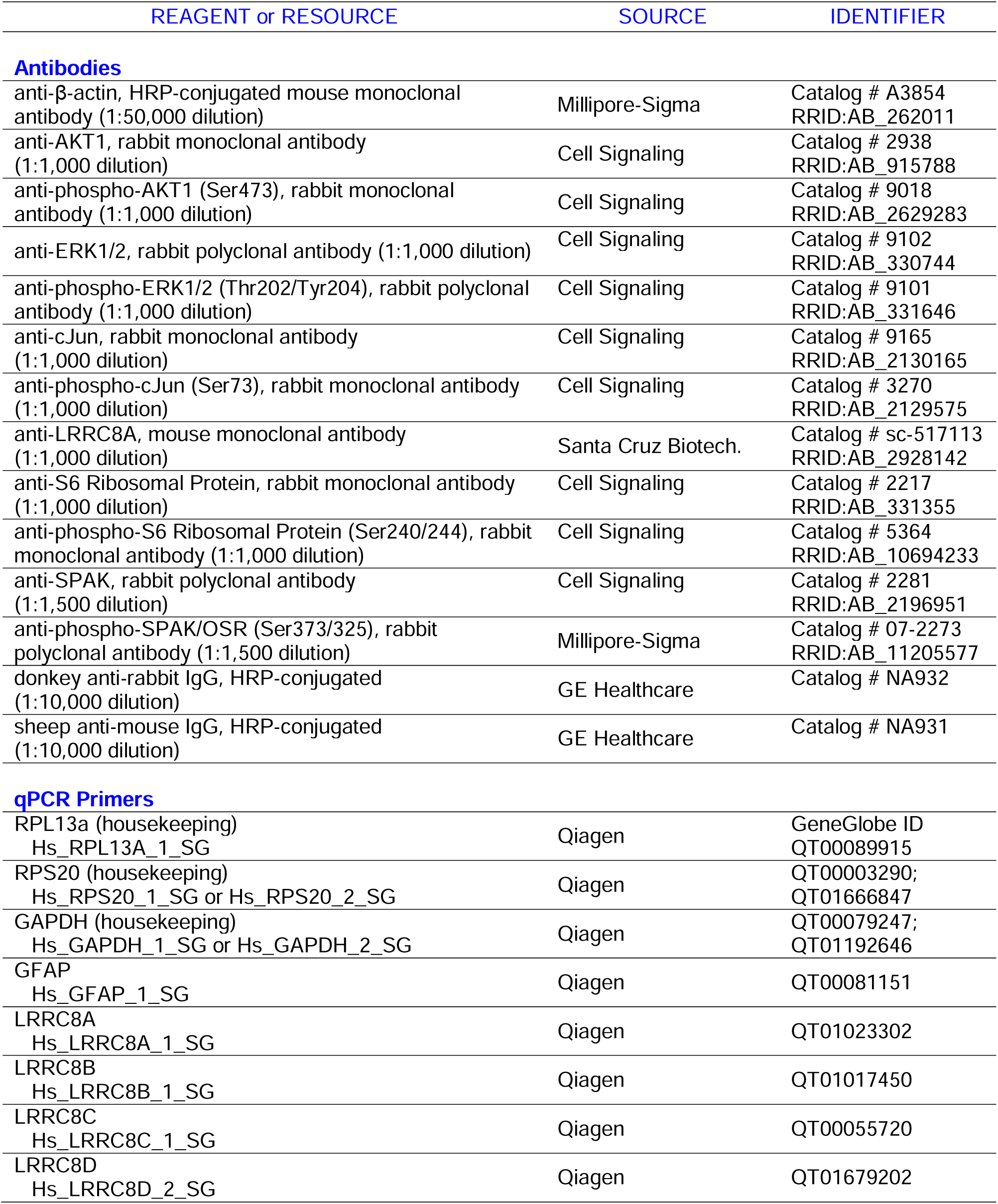
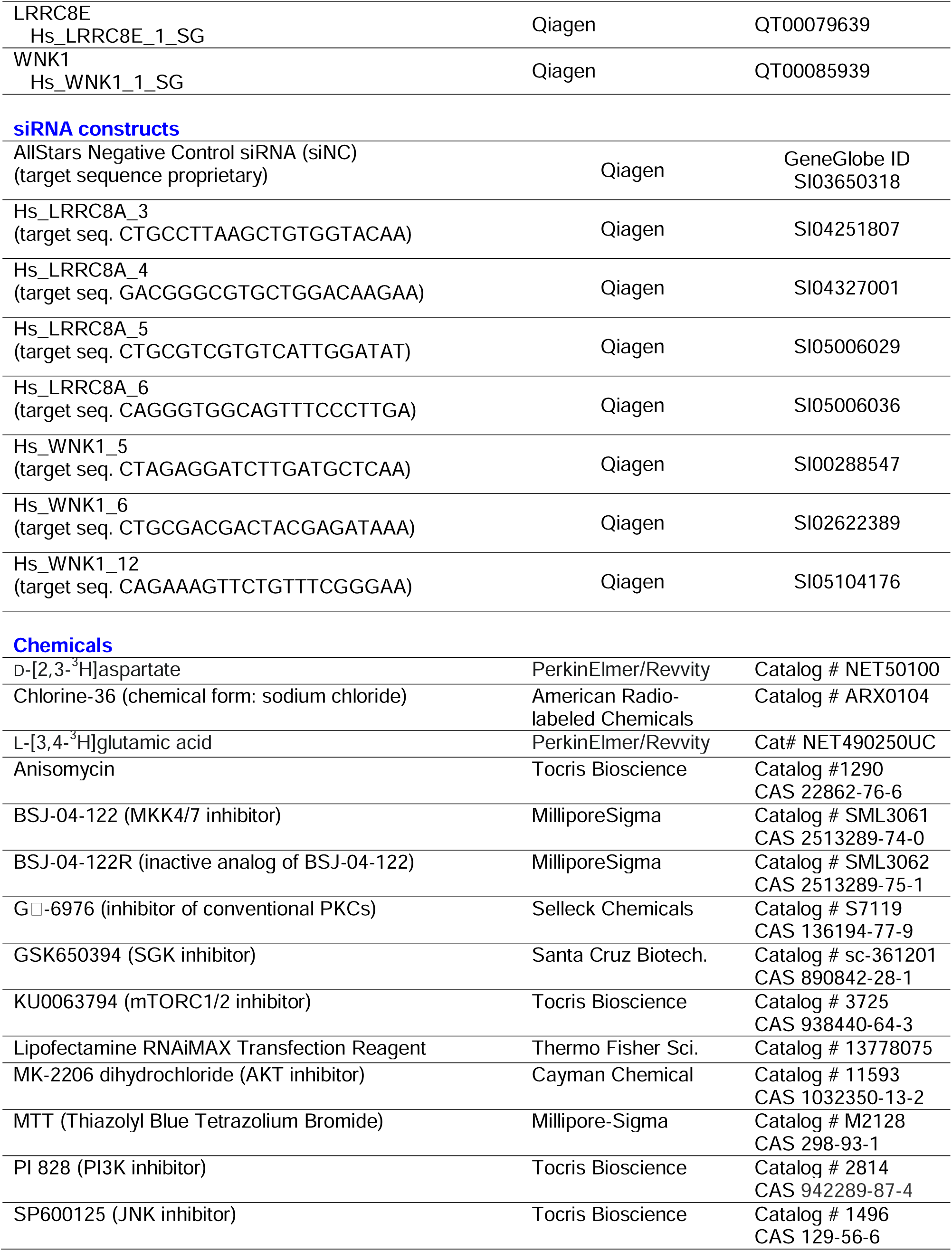
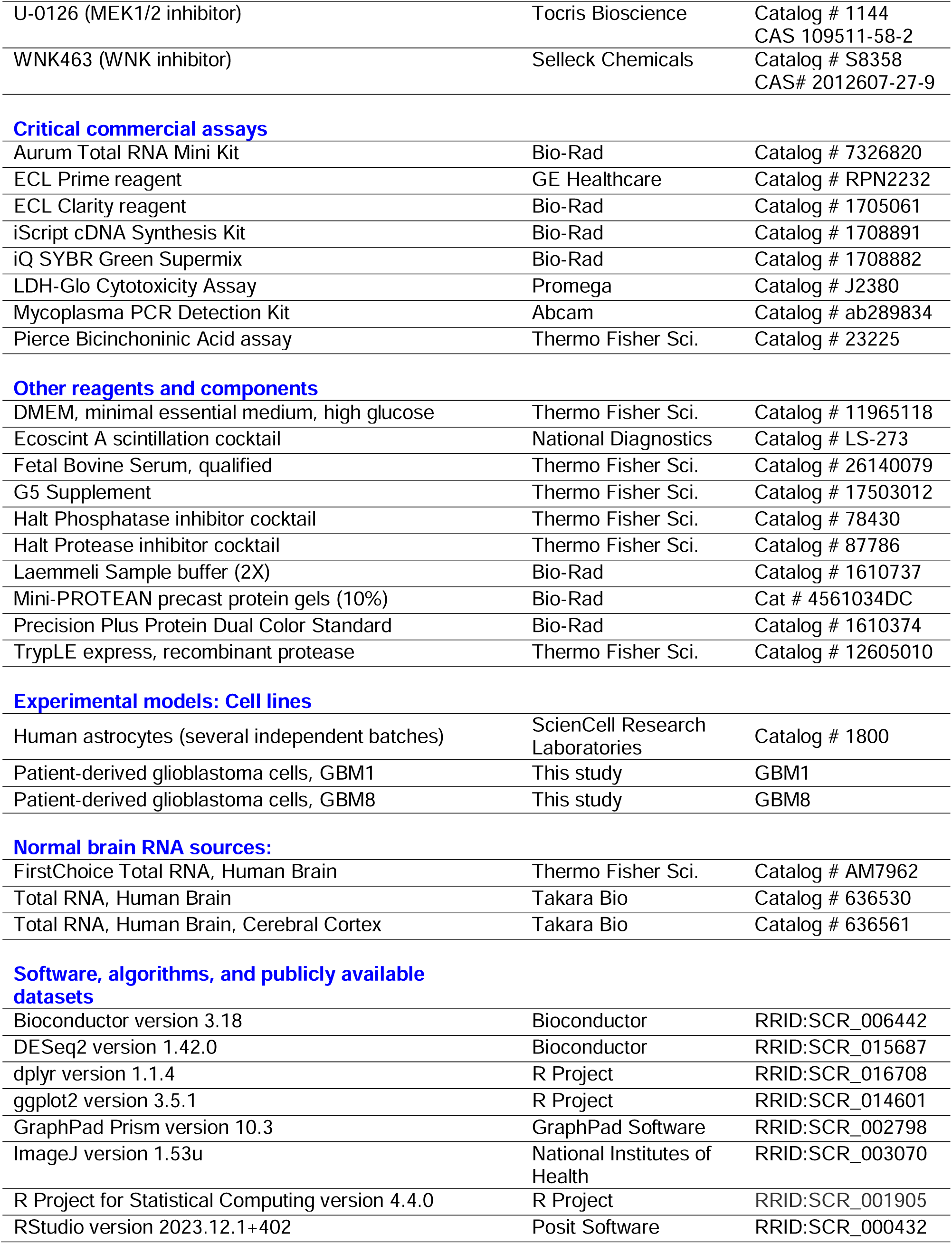
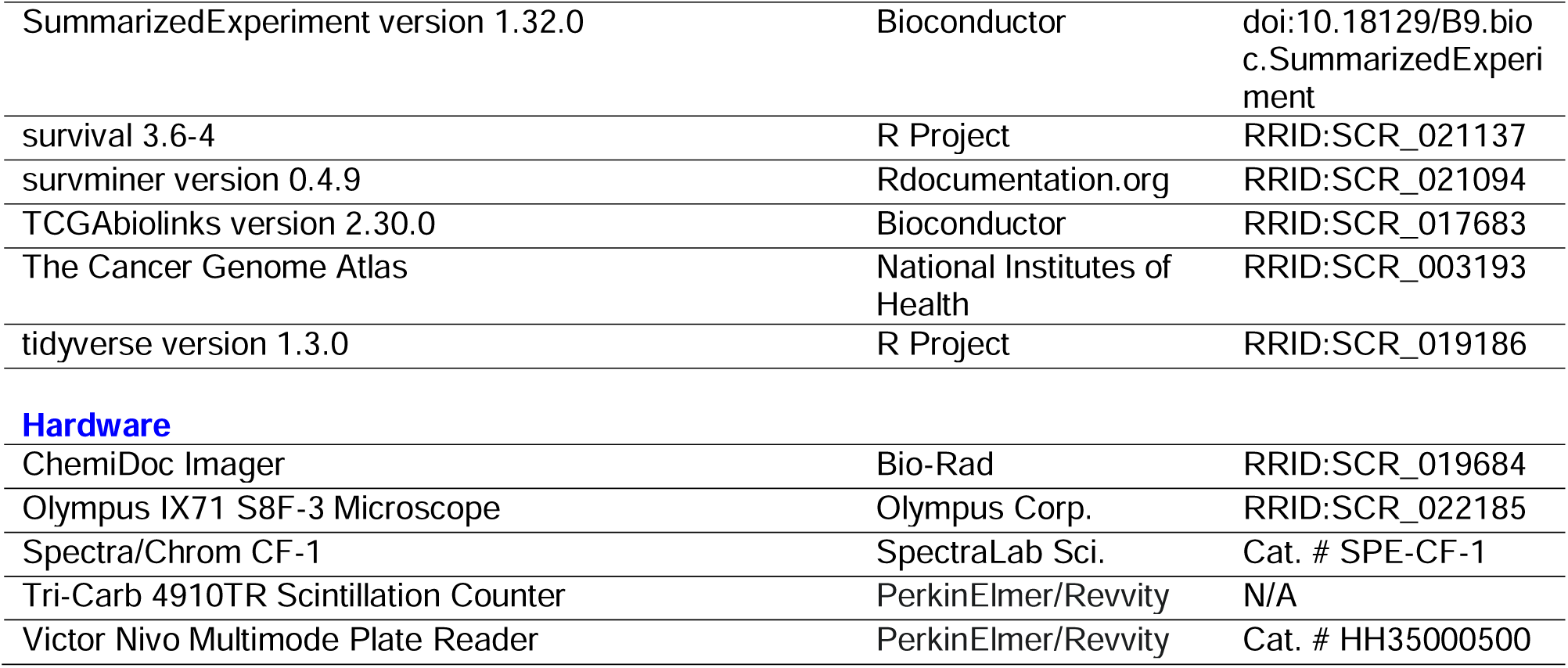

## RESOURCE AVAILABILITY

### Lead contact

Further information and requests for resources and reagents should be directed to and will be fulfilled (as available) by the Lead Contact, Alexander A. Mongin.

### Materials and availability statement

Due to the very limited amount of surgery derived GBM specimens, there are restrictions to the availability of GBM-derived cDNA and protein samples. Early passages of primary GBM cells are not available due to their scarcity. This study has used and validated unique molecular biology tools that are fully described in Key Resources Table and available from commercial sources. All primary data derived from this work are freely available as specified below.

### Data and code availability

- All patient samples used in this study have been deidentified
- All primary data used for analyses in this work are deposited in the Open Science Framework (osf.io) and are freely accessible for re-analysis at https://osf.io/2r7k9
- Original R codes used in the analysis of TCGA materials are deposited in the Open Science Framework and accessible at https://osf.io/sfvjg/ (TPM expression analysis) and https://osf.io/q6mvz (DEseq2-based patient survival analysis).

## EXPERIMENTAL MODEL AND STUDY PARTICIPANTS DETAILS

### Patient samples

Human GBM specimens were collected from 28 patients diagnosed with glioblastoma (WHO grade IV) and operated in the Department of Neurosurgery at Albany Medical Center System Hospital (AMCSH), Albany, New York, USA. The glioblastoma origin of tissue samples was confirmed during surgical resection by the standard histopathology in the Department of Pathology at AMCSH. The unused GBM tissue samples were collected in a sterile container filled with ice-cold saline, deidentified, and transferred on ice to further processing in a molecular biology laboratory. There, specimens were dissected from heavily vascularized parts by a sharp scalpel and separated into three parts for isolation of cDNA, generation of protein lysates, and preparation of primary glioblastoma cells as described below. All relevant procedures were approved by the Institutional Review Board of Albany Medical College (IRB # 2536, 6006). Written informed consent was obtained from the patients or their authorized representatives in accordance with the Declaration of Helsinki^115^ (rev.2024 https://www.wma.net/policies-post/wma-declaration-of-helsinki/, last accessed on 11/11/2024).

### Patient-derived primary GBM cell lines

GBM tissue specimens were washed in an ice-cold sterile phosphate-buffered saline (PBS), transferred into a Petri dish and cut into smaller fragments with a scalpel. Dissected tissue was further digested in the solution of 0.125% trypsin plus 0.015% EDTA for 10 minutes at 37°C. Cells were dissociated by mechanical trituration using a fire-polished glass Pasteur pipette, and the resulting suspension was filtered through a 70-μm cell strainer (Corning). Protease activity was quenched by adding an excess of Optimized Minima Essential Medium (Opti-MEM, Thermo Fisher Scientific) which was supplemented with 10% fetal bovine serum (FBS, Thermo Fisher Scientific), 100 U/ml penicillin and 100 µg/ml streptomycin (pen-step, Thermo Fisher Scientific). The final cell suspension was plated in a 75 cm^2^ flask (T75, Corning) and cultivated in a CO_2_ incubator in a mixture of 5% CO_2_/balance air at 37°C. After cells adhered to plastic, medium was replaced with high glucose Dulbecco’s Modified Eagle’s Medium (DMEM, Thermo Fisher Scientific) supplemented with 10% FBS and pen-step. Medium was fully replaced twice a week and GBM cell cultures were grown to ∼90% confluency. For propagation, cells were removed from a substrate using the recombinant protease TrypLE (Thermo Fisher Scientific) and replated into T75 or cryoprotected. Initial gene expression analyses were performed in passages 1 or 2. For all other described in this study experiments, GBM cells were used before passage 10 (in majority of cases at passages 4-6).

### Human GBM datasets in The Cancer Genome Atlas (TCGA)

We downloaded from TCGA (https://www.cancer.gov/ccg/research/genome-sequencing/tcga) all publicly available GBM datasets matching the following criteria: (i) identified as GBM, non-mutant (wild-type) isocitrate dehydrogenase (IDH1/2), (ii) complete RNAseq available, (iii) survival data available. Altogether, we analyzed data from 126 patients of both sexes (see Results section and additional information below). The healthy brain controls were sourced from the Gene Expression Omnibus repository and represent 29 brains (GEO series GSE196695).

## METHOD DETAILS

### Computational analysis of glioblastoma databases in TCGA

Publicly available GBM datasets were downloaded from TCGA and processed using an open-source programming language R (version 4.3.1), the R Studio platform (v. 2023.12.1+402), and the R package TCGAbiolinks (v. 2.30.0). The R codes used in our analyses have been deposited in the Open Science Framework, as referenced in the *Data and Code Availability* section. For comparisons of relative expression of multiple LRRC8 genes (LRRC8A-E) across GBM and normal brain tissue specimens, RNAseq data were downloaded and analyzed as transcript per million (TPM) counts. For additional survival analysis, patient LRRC8A RNAseq levels were downloaded as unstranded counts and normalized using the DEseq2 variance-stabilizing transformation. The latter normalization is considered superior to TPM and FPKM for comparisons of individual gene expression because it accounts for sample-to-sample differences in library size, sequencing depth, and expression profile.^59,60^ For survival analysis patients were separated into four quartiles and statistically compared using the Kaplan-Meier method (see *Statistical analyses*).

### mRNA isolation and real-time quantitative PCR

mRNA was extracted from tumor tissue or GBM cultures using the Aurum Total RNA Mini Kit (Bio-Rad Laboratories). Briefly, 100-300 mg of tissue was carefully dissected in a Petri dish using scalpel and forceps pretreated in diethylpyrocarbonate (DEPC) and lysed using the lysis buffer provided with a kit. In cell preparations, GBM cells were additionally washed from culture medium with Ca^2+^/Mg^2+^-containing PBS prior lysis. Lysates were homogenized by pipetting, mixed with equal parts ice-cold 70% EtOH and processed per manufacturer’s protocol. RNA concentrations were measured using the NanoDrop 1000 spectrophotometer (Thermo Fisher Scientific). Subsequent cDNA generation was done using the iScript cDNA synthesis kit (Bio-Rad Laboratories) using one µg of RNA per each 20 µLs of reaction volume. Resulting cDNA samples were aliquoted and stored at −80°C. mRNA expression levels were assessed by quantitative real-time PCR (qRT-PCR) in a CFX96 Real Time System (Bio-Rad) with validated quantitative PCR primers (Qiagen, see the Key Resourses Table) and SYBR Green Master Mix (Bio-Rad). Expression levels were calculated by the ΔCt (for tissue and cell expression) or ΔΔCt method^116^ (in RNAi analyses). Data were normalized within the sample to the housekeeping gene RPL13a. GAPDH and RPS20 were probed as additional housekeeping and quality controls.

### Protein isolation and Western blotting analysis

Proteins were extracted from tumor tissue samples or cell cultures by lysis in the solution of 2% SDS plus 8 mM EDTA, additionally containing Halt phosphatase and protease inhibitors (Thermo Fisher Scientific). A small aliquot of each lysate was used for determining protein content using the Pierce Bicinchoninic acid protein assay kit (Thermo Fisher Scientific). The remaining lysates were mixed with 2× reducing Laemmli buffer (Bio-Rad), aliquoted, and stored at −80°C until analysis. 20-µg protein samples were loaded on precast 10% acrylamide gels (Bio-Rad), separated using SDS-PAGE, and subsequently transferred onto PVDF membrane (Bio-Rad) using a Trans-Blot Turbo transfer system (Bio-Rad). Membranes were next blocked for 15 min in either 5% fat-free milk or EveryBlot Blocking Buffer (Bio-Rad, used for all phosphoprotein analyses) and incubated with primary antibodies overnight at 4°C. After several washes, membranes were additionally incubated with an appropriate HRP-conjugated secondary antibody for 90-min at room temperature. Chemiluminescence signal of proteins of interest was captured using the ChemiDoc MP Imaging System (Bio-Rad) and quantified using an ImageJ software^117^ (National Institute of Health, v. 1.53u). To ensure equal protein load, one-to-two days after initial western blotting, membranes were re-probed without stripping with the HRP-conjugated antibody recognizing the housekeeping protein β-actin. Protein levels were normalized to housekeeping protein signal, and further double normalized to average immunofluorescence in at least two ‘control’ samples on the same Western blot membrane (e.g., ‘control siRNA’ or ‘vehicle treated’). In assays measuring phosphospecific protein signal, immunoluminescence values were additionally analyzed by dividing normalized phosphoprotein signal by normalized total protein signal in the same lysate. (all antibodies and their dilution factors are listed in *Key Resources Table*).

### siRNA manipulation of gene expression

siRNA transfections were carried out using Lipofectamine RNAiMAX reagent (Thermo Fisher Scientific) following the manufacturer’s recommendations. In short, siRNA/Lipofectamine RNAiMAX complexes were prepared in Opti-MEM, in a light-protected environment, and then added to cell cultures bathed in a serum-free Opti-MEM. The final siRNA concentration was 10, 25, of 50 nM (optimized for each construct). After an initial 4-h incubation, Opti-MEM was supplemented with DMEM containing 10% FBS and 0.5% penicillin-streptomycin. The efficacy of siRNA constructs was measured 48-h after transfection using qRT-PCR (see the Key Resources Table for the list of siRNA constructs and target sequences). Protein isolation and functional assays were performed 96-hours after transfection.

### Radiotracer assays of VRAC activity

Activity of VRAC was quantified by measuring the release of the radiolabeled VRAC-permeable, non-metabolizable analog of glutamate D-[^3^H]aspartate. GBM cells or primary human astrocytes were grown to confluency on 18-mm square coverslips and subsequently treated with siRNA as indicated. After 72 h, cells were additionally incubated overnight with 2 μCi/ml D-[^3^H]aspartate (PerkinElmer) in DMEM+10% FBS in a CO_2_ incubator at 37°C. The next day, cells were washed from intracellular isotope with warm chemically defined isoosmotic Basal medium containing (in mM) 135 NaCl, 3.8 KCl, 1.2 MgSO_4_, 1.3 CaCl_2_ mM, 1.2 KH_2_PO_4_, 10 D-glucose, and 10 HEPES (pH 7.4, osmolarity 290 ±3 mOsm). Coverslips were loaded in a Lucite perfusion chamber with Teflon top that leaves approximately 200-300 µm above coverslip, and superfused at the rate of ∼1.2 ml/min with either isoosmotic Basal medium or hypoosmotic medium. The composition of hypoosmotic medium was similar with the exception that NaCl concentration was reduced to 85 mM leading to a 30% decrease in medium osmolarity (190 ±3 mOsm, verified by a freezing point micro-osmometer µOsmette, Precision Systems, Natick, MA, USA). One-minute perfusate fractions were collected using an automated fraction collector (Spectra/Chrom CF-1, SpectraLab, Markham, ON, Canada). Individual superfusate fractions were next mixed with an Ecoscint A scintillation cocktail (National Diagnostics, Atlanta, GA, USA), and the levels of released D-[^3^H]aspartate in each sample were determined using a Tri-Carb 4910TR scintillation counter (PerkinElmer). To determine the amount of radiotracer remaining inside cells on a coverslip, after completion of experiment, cells were lysed in the solution of 2% SDS plus 8mM EDTA. Fractional isotope release rates were calculated using an Excel-based custom algorithm.

### Measurements of intracellular Cl levels

Intracellular chloride levels were analyzed by measuring steady-state accumulation of the radiotracer ^36^Cl^−^. Cells were grown to confluency in six-well plates, in DMEM+10% FBS in the atmosphere containing 5% CO_2_/balance air at 37°C. Confluent cells were equilibrated overnight in Opti-MEM +10% FBS. To initiate isotope uptake, an aliquot of Opti-MEM+FBS additionally containing ^36^Cl^−^ (American Radiolabeled Chemicals) was added to each well to the final isotope activity 0.5 µCi/ml. After 20-min incubation, isotope uptake was terminated by washing four times with an ice-cold solution containing (in mM) 300 D-mannitol, 1.2 MgSO_4_, and 10 HEPES (pH 7.4, 320 ± 3 mOsm). After the final wash, cells were lysed in 2% SDS plus 8 mM EDTA. Cell lysates were mixed with the scintillation cocktail Ecoscint A (National Diagnostics) and intracellular ^36^Cl^−^ levels were determined using a Tri-Carb 4910TR scintillation counter (Revvity). ^36^Cl^−^ radioactivity was further normalized to protein levels in each well.

### Cell proliferation assays

Cellular proliferation was measured using the MTT proliferation assay. Cells were plated at low densities (15K/well for GBM1 or 30K/well for GBM8) and allowed to adhere overnight. Cells were then transfected with siRNA (see above) or treated with pharmacological inhibitors as indicated in figure legends. After 4 days of transfection or 3 days of pharmacological treatments, cell culture media were washed with isoosmotic Basal medium and incubated in the Basal medium containing 0.5 mg/ml MTT for 1-2 hours at 37°C. MTT-containing media were removed, and the newly formed intracellular formazan crystals were solubilized with dimethyl sulfoxide (DMSO). The optical density of the formazan solutions were determined by measuring the absorbance at 562 nm in an ELx800 plate reader (BioTek). Data are presented as the percentage of MTT signal compared to within-plate untreated or negative control cells.

In addition, DAPI-stained nuclei counting was used as an alternative method to ascertain the effect of certain treatments on cell proliferation. Cells were plated and treated as described above. On the final day, cell culture medium was aspirated, and cells were fixed in 4% paraformaldehyde in PBS for 15 minutes at room temperature (22-23°C). 4% PFA/PBS solution was aspirated, wells were gently washed twice with cold Ca^2+^/Mg^2+^-free DPBS and incubated in ice-cold DPBS containing 300 nM DAPI for 20 minutes at room temperature in a light-protected environment. DAPI-stained nuclei were imaged at 10× magnification using a Keyence BZ-X800 fluorescent microscope (Keyence Corporation) and counted using ImageJ.

### Fluorescence-assisted cell sorting (FACS) analysis of cell cycle stage

Cellular DNA content was determined by staining cells with propidium iodide and measuring fluorescence using a Becton Dickinson FACScan (Rutherford, NJ, USA). Cells were harvested by trypsinization and fixed in cool 70% ethanol for 6 h. Subsequently, fixed cells were incubated in a solution containing 1 mg/ml RNase and 20 μg/ml propidium iodide for 30 min. For each cell population, 10,000 cells were analyzed by FACS and the proportion of cells in G_0_/G_1_, S, and G_2_/M phases was measured estimated using the FlowJo program (version 10.0).

### Cell death assay

Cell death was assessed by measuring levels of lactate dehydrogenase (LDH) release using a luciferase reporter-based LDH cytotoxicity assay (LDH-Glo, Promega Corporation). To reduce contribution of nonspecific LDH signal originating from FBS, cells were grown in Opti-MEM plus G5 serum-free supplement. After 1, 3, or 4 days, cell culture media from each well was sampled, mixed with equal parts of LDH storage buffer, and transferred onto ice. The remaining cells were lysed in LDH lysis buffer. Cell lysates were further diluted in an LDH storage buffer and kept on ice. Supernatants and cell lysates were transferred into 96-well plates, mixed with a freshly prepared detection reagent, and levels of luminosity were measured using a Victor Nivo 3T Multimode Plate Reader (Revvity). As a positive control for cell death, cells were treated with the protein kinase inhibitor staurosporine (STS; 300nM, 24-hours prior to processing). Relative LDH release levels were calculated by dividing the background-corrected LDH signal in supernatants by the total LDH levels in cell lysates from the same wells. Cell death values are reported as percentages.

### QUANTIFICATION AND STATISTICAL ANALYSIS

All data are presented as the mean values ±SD, with the number of independent experiments (n) indicated in each figure. n represents an independently prepared and treated cell culture, or an individual patient in human datasets. Experimental data were tested for normal distribution. In most cases, statistical differences between groups with normal distribution were determined by one-way ANOVA with Dunnett’s post hoc correction for multiple comparisons. The two-group comparisons were done using an unpaired t-test. In those instances where data were normalized to within-experiment controls, statistical comparisons were performed using a one-sample t-test with the Bonferroni correction. Gene expression data were compared using an unpaired t-test with the Welch correction and Holm-Šídák adjustment. The survival rates for GBM patients were compared using the Gehan-Breslow-Wilcoxon test. Plotting and statistical analyses were done using Prism software (version 10.3.0, GraphPad Software).

