## Supplemental Figures for "LRRC8A-containing anion channels promote glioblastoma proliferation via a WNK1/mTORC2-dependent mechanism"

**Supplemental Figure 1 • Fidaleo, Bach *et al.***

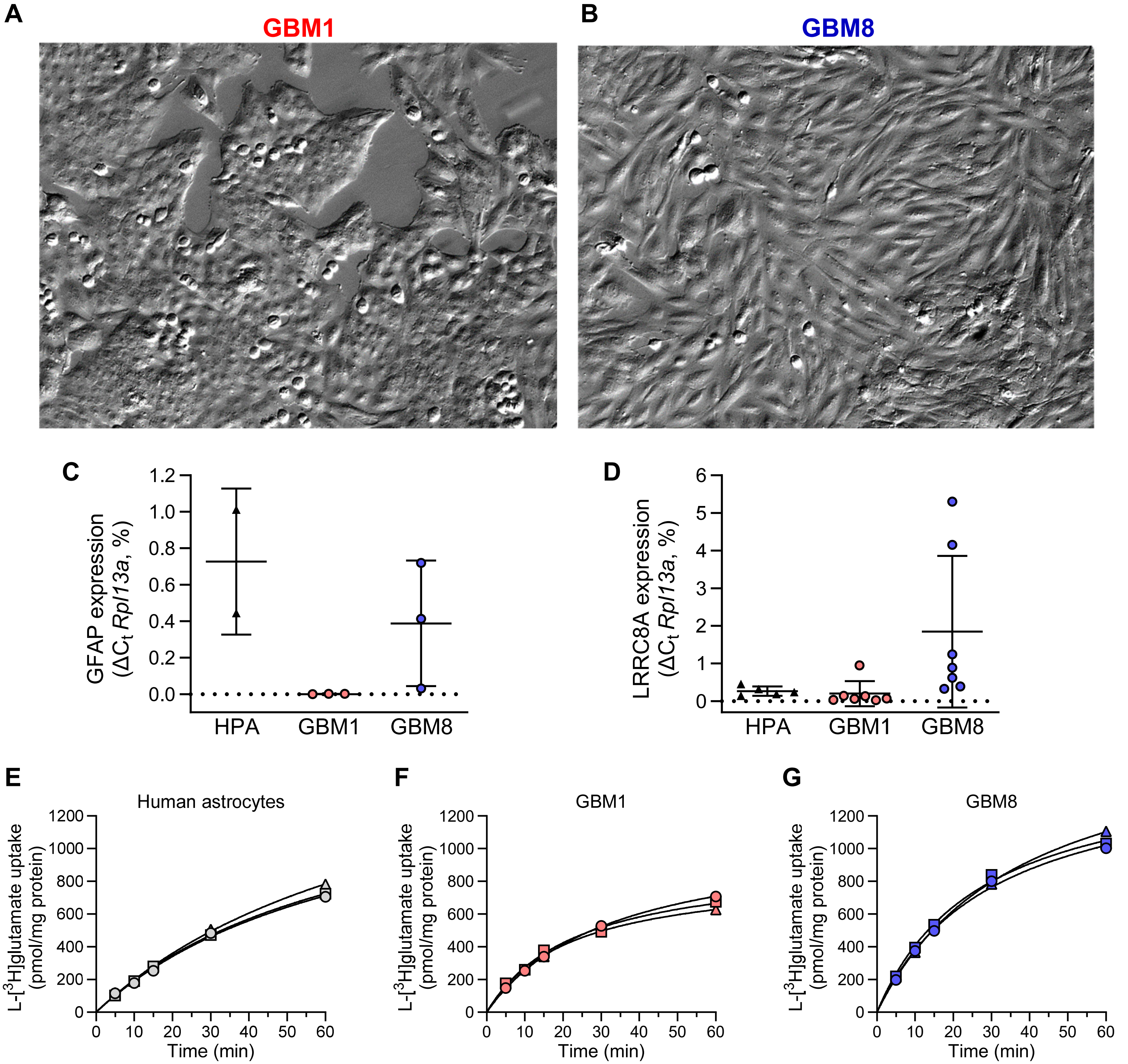


### Properties of the patient-derived glioblastoma cell lines, GBM1 and GBM8

(**A-B**) Representative images of the patient-derived GBM1 (A) and GBM8 (B) cells used in Figure 1B but shown without cropping. Images were acquired with an Olympus IX71 microscope using Hoffman optics and the identical 160× magnification

(**C**) Comparison of *Gfap* mRNA expression levels in patient derived GBM cell cultures (n=3) and primary human astrocytes (n=2).

(**D**) Comparison of *Lrrc8a* mRNA expression levels in patient derived GBM cell cultures (n=7) and primary human astrocytes (n=5).

(**E-G**) Kinetics of L-[$^3$H]glutamate uptake in human astrocytes (E), GBM1 cells (F), and GBM8 cells (G). Data are the mean values ± SD of 3 separate wells from 3 independent assays.

**Supplemental Figure 2 • Fidaleo, Bach *et al.***

A

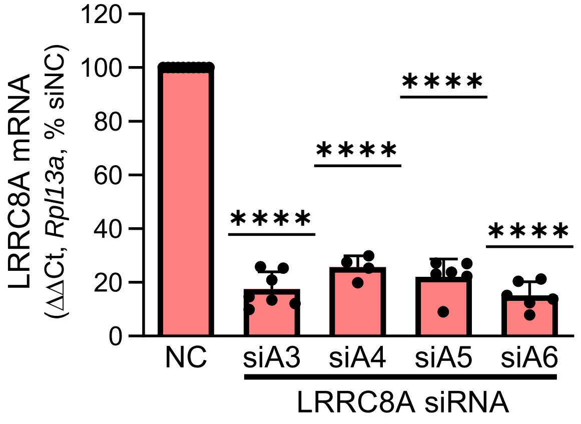


B

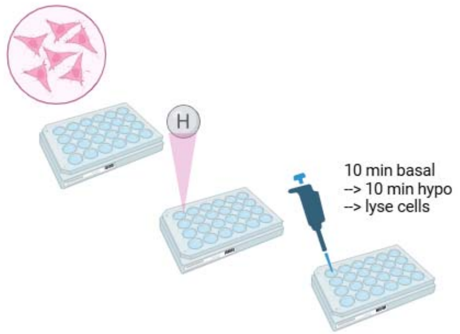


C

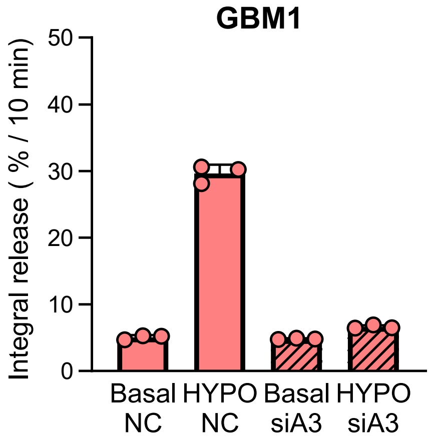


D

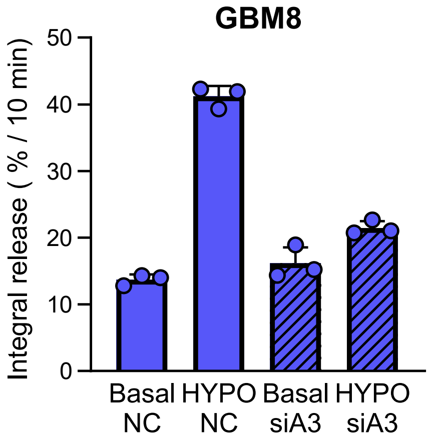


### Validation of siRNA-based knockdown of LRRC8A and VRAC

(**A**) Effect of LRRC8A siRNA on *Lrrc8a* mRNA levels. GBM1 cells were transfected with the LRRC8A-specific siRNAs siA3, siA4, siA5, and siA6, or the negative control siNC. Expression values were measured using qRT-PCR and normalized to *Rpl13a.* Data are the mean values ±SD. n=4-7 independent transfections. One-sample t-test with Bonferroni correction, ****p<0.0001, vs. siNC.

(**B**) Schematic depicting workflow for experiments measuring VRAC activity as swelling-activated release of D-[$^3$H]aspartate. Cells were plated to 24-well plates, loaded overnight with D-[$^3$H]aspartate, washed from extracellular radiotracer, and then exposed for 10 min to isoosmotic (Basal) and 10 min hypoosmotic (HYPO, -30% osmolarity) media. Extracellular media were collected, and cells were lysed to determine the remaining radiotracer content. Data are presented as the integral 10-min release values normalized to the D-[$^3$H]aspartate content at the beginning of incubation.

(**C**) Functional downregulation of VRAC activity in GBM1 cells treated with LRRC8A siRNA (siA3) as compared to the negative control siNC. Data are the mean values ±SD. n=3 independent transfections.

(**D**) Functional downregulation of VRAC activity in GBM8 cells treated with LRRC8A siRNA (siA3) and compared to the negative control siNC. Data are the mean values ±SD. n=3 transfections.

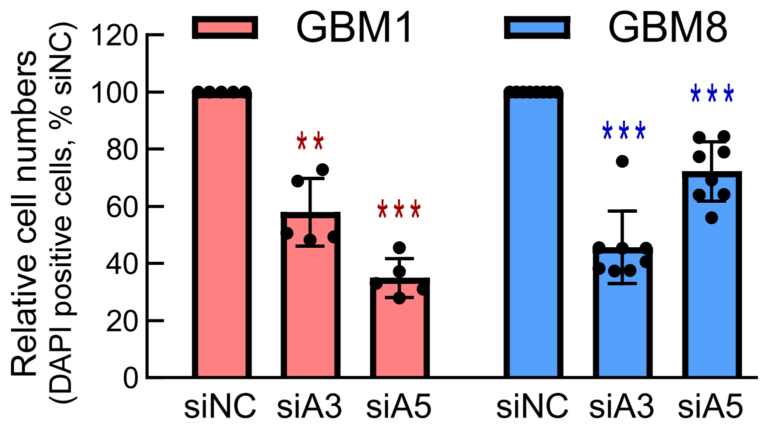


**Validation of the anti-proliferative effect of LRRC8A knockdown in GBM1 and GBM8 cells using automated counts of DAPI-stained cells**

GBM1 and GBM8 cells were plated into 24-well plates at 10,000 (GBM1) or 20,000 (GBM8) cells/well, transfected with the LRRC8A-specific siRNAs (siA3 and siA5), or the negative control siRNA (siNC). 96 h post-transfection, cells were fixed in 4% paraformaldehyde, permeabilized with 0.1% Triton X-110, and stained with DAPI (1 ng/ml). DAPI-stained nuclei were counted using a Citation plate reader and Gen5 software (BioTek Instruments). Cell numbers were additionally normalized to the siNC-treated controls in each expariment. Data are the mean values ±SD. n=5-8 independent transfections/cell line. **p<0.01, ***p<0.001, vs. siNC. One-sample t-test with Bonferroni correction for multiple comparisons.

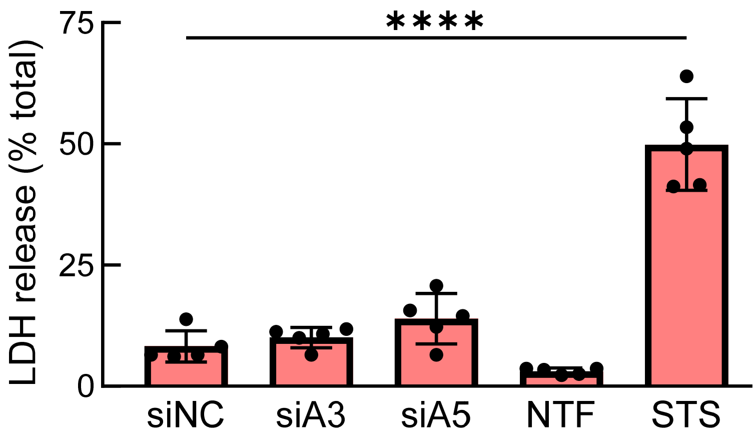


**LRRC8A knockdown does not increase cell death in GBM1 cultures**

GBM1 cells were plated into 24-well plates at the density of 20,000 cells/well, transfected with the LRRC8A-specific siRNAs (siA3 and siA5), or the negative control siRNA (siNC), and grown for additional 96 h. Cell death was quantified by measuring extracellular levels of the cytosolic enzyme lactate dehydrogenase (LDH) using a luciferin-luciferase-based LDH kit (Promega). The extracellular LDH levels were normalized to total LDH content in each well and presented as % of total LDH. As controls, we measured LDH release in non-transfected cells (NTF) and cells treated for 24 h with the apoptosis-inducing agent staurosporine (STS; 300 nM). Data are mean values ± SD. n=5 independent transfections or treatments. ****$p<0.0001$, vs. siNC, One-way ANOVA with Dunnett's post hoc correction for multiple comparisons. There were no statistical differences between cell cultures treated with siNC, siA3, and SiA5 constructs.

**Supplemental Figure 5 • Fidaleo, Bach *et al.***

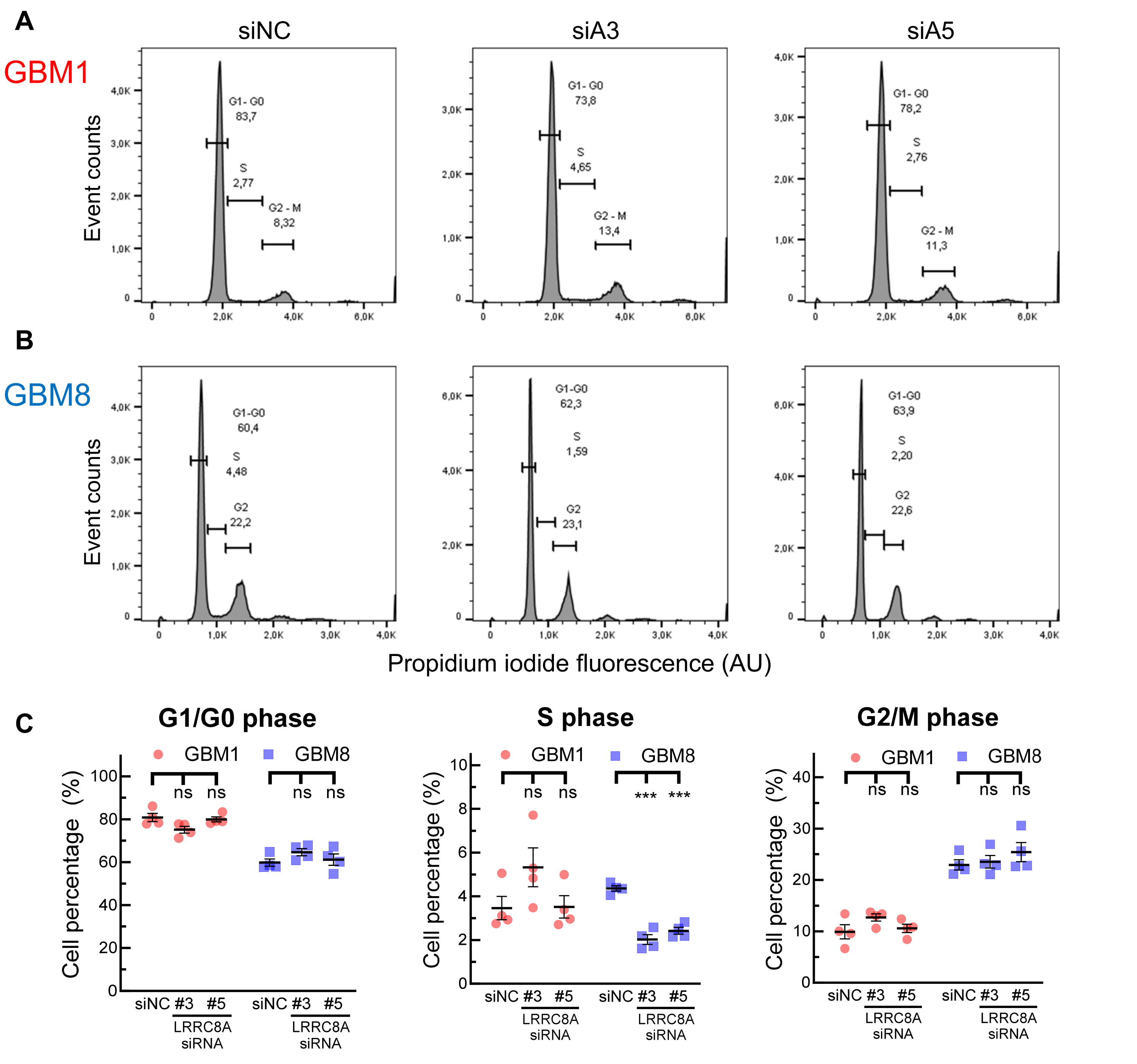


### Effect of LRRC8A downregulation on cell cycle distribution in GBM1 and GBM8 cells

GBM1 or GBM cells were transfected with the LRRC8A-specific siRNAs (siA3 and siA5) or the negative control siRNA (siNC). 96 h post-transfection, cells were detached from the substrate and stained with propidium iodide. Cell cycle distribution was measured using a fluorescence-activated cell sorting analysis (FACS). For additional methodological details see *Star★Methods*.

(**A-B**) Representative histograms of FACS analysis in GBM1 (A) and GBM8 (B) cells.

(**C**) Quantification of FACS analyses of cell cycle distribution across various phases ($G_1/G_0$, S and $G_2/M$). Data are the mean values ±SD. n=4 independent transfections. ***p<0.001, ns=not significant, vs. siNC. One-way ANOVA with Bonferroni correction.

**Supplemental Figure 6 • Fidaleo, Bach *et al.***

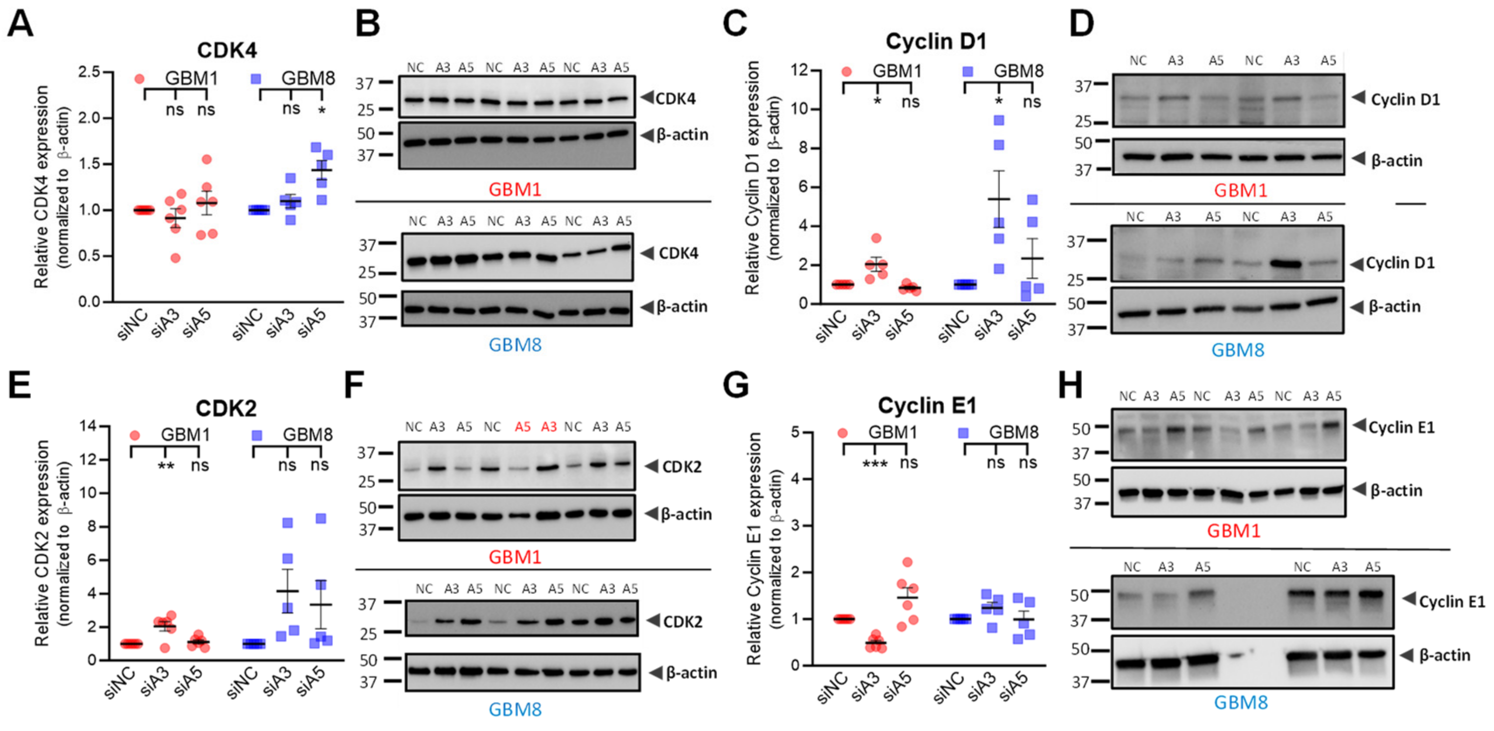


### Effect of LRRC8A knockdown on levels of cell cycle-related proteins in GBM1 and GBM8 cells

(**A-B**) Quantification (A) and representative images (B) of Western blot analysis of the effect of LRRC8A downregulation on CDK4 expression in GBM1 and GBM8 cells. Cells were transfected with the LRRC8A-specific siRNA (siA3 and siA5) or the negative control siRNA (siNC). n=5-6 independent transfections per cell line.

(**C-D**) Quantification (C) and representative images (D) of Western blot analysis of the effect of LRRC8A downregulation on cyclin D1 expression in GBM1 and GBM8 cells. Same treatments were performed as in panel A. n=5 independent transfections per cell line.

(**E-F**) Quantification (E) and representative images (F) of Western blot analysis of the effect of LRRC8A downregulation on CDK2 expression in GBM1 and GBM8 cells. Same treatments were performed as in panel A. n=5-6 independent transfections per cell line.

(**G-H**) Quantification (G) and representative images (H) of Western blot analysis of the effect of LRRC8A downregulation on cyclin E1 expression in GBM1 and GBM8 cells. Same treatments were performed as in panel A. n=5-6 independent transfections per cell line.

For A-H, All data points are normalized to the within-set negative control siNC and presented as individual points with mean ± SD. ns=$p>0.05$, *$p<0.05$, **$p<0.01$, ***$p<0.001$ vs. siNC. One-sample t-test with Bonferroni correction.

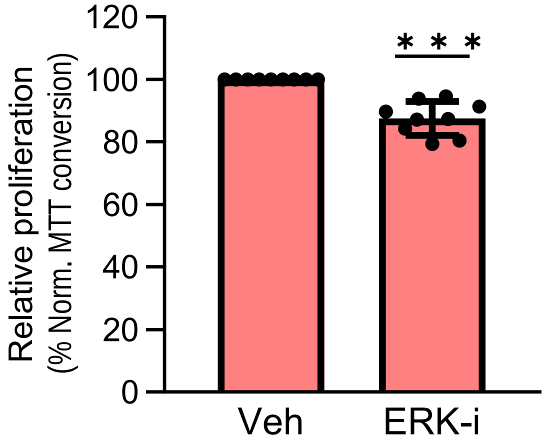


**Effect of inhibition of ERK signaling with trametinib on GBM1 proliferation**

GBM1 cells were plated in 24-well plates, treated with the MEK1/2 inhibitor Trametinib (Tram, 0.3 μM) or vehicle control (Veh, 0.1% DMSO), and grown for 72 h. Cell proliferation values were measured using the MTT assay and normalized to within-plate vehicle-treated controls (n=9). Data are the mean values ± SD. ***p<0.001, vs Veh. One-sample t-test.

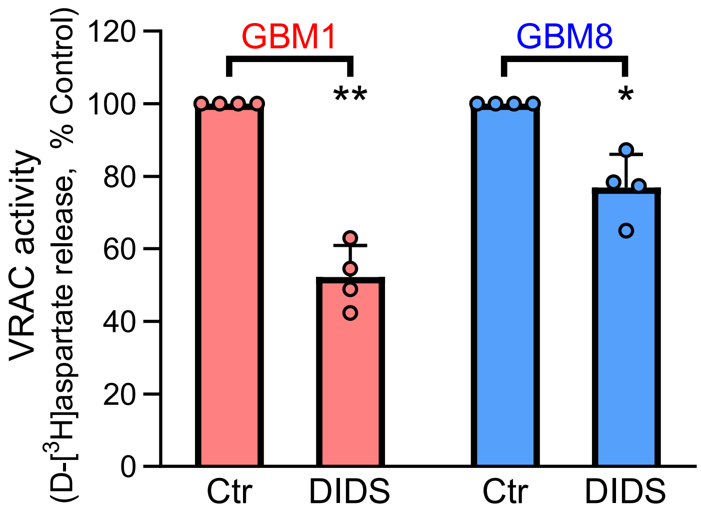


**Effect of the chloride channel blocker DIDS on VRAC activity in GBM1 and GBM8 cells measured in serum containing media.**

GBM1 or GBM8 cells were grown in 24-well plates and loaded overnight with D-[$^3$H]aspartate. On the day of the assay, extracellular radiotracer was washed out, and cells were exposed for 10 min to serum-containing cell culture medium in which osmolarity was maintained or reduced by 30% by adding water. VRAC activity was measured as swelling-activated release of D-[$^3$H]aspartate, in the presence or absence of 500 µM DIDS. VRAC activity values were normalized to within-experiment controls (Ctr). n=4/cell line. Data are the mean ± SD. ***$p<0.001$, **$p<0.01$, vs Control. One sample t-test.
